# Modified Potential Functions Result in Enhanced Predictions of a Protein Complex by All-Atom MD Simulations, Confirming a Step-wise Association Process for Native PPIs

**DOI:** 10.1101/241810

**Authors:** Zhen-lu Li, Matthias Buck

## Abstract

The relative prevalence of native Protein-Protein interactions (PPIs) are the cornerstone for understanding the structure, dynamics and mechanisms of function of protein complexes. In this study, we develop a scheme of scaling the protein-water interaction in the CHARMM36 force field, in order to better fit the solvation free energy of amino acids side chain analogues. We find that the molecular dynamics simulation with the scaled force field, CHARMM36s, as well as a recently released version-CHARMM36m, effectively improve on the overly sticky association of proteins, such as ubiquitin. We investigate the formation of a heterodimer protein complex between the SAM domains of the EphA2 receptor and SHIP2 enzyme by performing a combined total of 48 µs simulations with the different potential functions. While the native SAM heterodimer is only predicted at a low rate of 6.7% with the original CHARMM36 force field, the yield is increased to 16.7% with CHARMM36s, and to 18.3% with CHARMM36m. By analyzing the 25 native SAM complexes formed in the simulations, we find that their formation involves a pre-orientation guided by Coulomb interactions, consistent with an electrostatic steering mechanism. In 12 cases, the complex could directly transform to the native protein interaction surfaces with only small adjustments in domain orientation. In the other 13 cases, modest orientational and/or translational adjustments are needed to reach the native complex. Although the tendency for non-native complexes to dissociate has nearly doubled with the modified potential functions, a dissociation followed by a re-association to the correct complex structure is still rare. Instead the remaining non-native complexes are undergoing configurational changes/surface searching which, however, rarely lead to native structures on a timescale of 250 ns. These observations provide a rich picture of the mechanisms of protein-protein complex formation, and suggest that computational predictions of native complex protein-protein interactions could be improved further.

## INTRODUCTION

Biological function that arises from the formation (and/or dissociation) of protein complexes critically depends on the strength and kinetics of the underlying protein-protein interactions (PPIs), where specific/native interactions typically compete with less specific/non-native contacts. Our detailed knowledge of PPIs and understanding of protein complex formation is far from complete.^1–5^ Especially, an appreciation of the protein association processes involved is still lacking for the great majority of systems at the molecular level. This is in part due to the limited temporal, if not spatial resolution of experimental methods. Despite early successes in Brownian and coarse grained molecular dynamics simulations of (mostly tightly bound) complexes,^2,6–10^ there have, as yet, been very few simulations of protein association processes using unrestrained all-atom simulations.^11–16^ In many cases it remains difficult to computationally predict the correct protein complexes structures ab initio. ^17^ Only relatively recently have computational protein docking programs started to incorporate main chain (or even side chain) flexibility;^18–19^ also the role of local, as well as global dynamics in protein complexes and in complex formation is now becoming well appreciated.^20–23^ Indeed, a community wide challenge, the critical assessment of predicted interactions (CAPRI), established in 2001, has yielded steady improvements over the years; currently most “easy to moderately difficult” protein complex target structures can be predicted reliably to be among the top 5-10 of calculated solutions,^17,24^ although often sequence/evolution-based as well as experimental information are used as restraints.

Classical all-atom molecular dynamics simulations are purely based on empirically derived physical potential functions and are typically conducted under one set of conditions. In principle, these simulations can provide information on the unrestrained and dynamic processes of protein association/dissociation as well as on the physical-chemical details of the interactions, which are not easily accessible by other computational methods or by current experiments. For example, recently, we reported all-atom simulations on the dissociation process of a small protein heterodimer, the EphA2-SHIP2 SAM: SAM domain complex.^25^ In that study we showed that a simple (e.g. a single) dissociation pathway was absent and that instead a step-wise process was involved, allowing for multiple pathways.^25^ A number of inferences could be made from that study, including the suggestion that the reverse process, that of protein association, would involve transient encounter complexes which may proceed to the native complex via intermediary bound states. Another process associated with protein complex formation-electrostatic steering-which directs the formation of encounter complexes to be close to the native protein configuration, has been reported and discussed over several decades.^26–28^

It is, however, still very challenging to accurately predict the native protein-protein interactions *ab initio* using classical all-atom molecular dynamics simulation. Insufficient sampling is a major obstacle in the accurate prediction of protein complexes but so are inaccuracies in the potential function. Once two proteins are trapped into a local minimum/non-native complex, they find it hard to dissociate or to convert to a native complex, as the relaxation time of dissociation or orientational transitions could be far beyond the simulation time scale. This could be the case with our system as well: here, using CHARMM36, the SAM domains associated on a reasonable timescale, but the initial associated states were often “trapped”, in that they showed few interconversions and fewer dissociation events, than would be expected from initial non-native encounter complexes.^9,29,30^ Consequently, the yield rates of native protein complexes were typically low within the available simulation time. Even with µs simulations of the native structure we found a lack of transitions between certain configurational states^22^. In fact, a low yield of correct structures, approx.7-8% is actually similar to those seen in the early Brownian Dynamics simulations^.9,10^ Because of this “kinetic trapping issue”, the molecular dynamics community has recently appreciated that several of the current potential functions appear to lack accuracy, especially for simulations involving protein association and for the behavior of intrinsically denatured proteins (IDPs).^14,31^ In case of the former, many proteins-even those having only non-specific non-covalent self-association (such as ubiquitin)-showed a strong and unphysical aggregation.^14,33^ Similarly, IDPs became too compact in the simulations.^33,34^ Both effects arise from an underestimation of protein solvation and both are in disagreement with experimental observations.

To address these problems, researchers have tried to improve parameters in the potential functions in different ways. In one study, Piana et al. developed a new water model, which could better reproduce intrinsically disordered proteins,^34^ and this has recently been refined to also work on folded proteins.^35^ A most recently released version of the CHARMM potential function, CHARMM36m, improves the prediction of α-helices and intrinsic disorder in polypeptides.^36^ Best et al. have explored a scaling of protein-water interactions with the Amber potential function,^37^ which showed an improved prediction for IDPs and non-specific protein associations. A study of Feig and colleagues found that the protein translational and especially rotational diffusion slowed down following an adjustment of the potential function by globally scaling protein-water interactions.^14^ Several other studies have rescaled the protein-protein interactions or protein-water interactions, or carried out a mixed rescaling with different water models (such as TIP4P) in order to get a better performance in the prediction of native protein configurations or protein-protein association.^38–42^

At this time, there are still only a few studies of unrestrained protein-protein association involving binding partners with moderate binding affinity using classical molecular dynamics simulations. Furthermore, it was not clear to what extent the potential, such as the recently revised version of CHAMM36, CHARMM36m, may accurately predict the native protein-protein interactions. Here we tested two model systems: the self-association of ubiquitin and the hetero-association between EphA2 SAM (E-SAM) domain and SHIP2 SAM (S-SAM) domain. The ubiquitin has very weak non-specific association with a dissociation constant K_d_ about 4.8 mM; while SAM domains have a moderate binding affinity with a K_d_ value at 2.2 µM, a value typical for the interaction of cell signaling proteins. In this study, we first applied a scheme of scaling the protein-water interaction to the original CHARMM36 potential function, thus it could better fit the solvation free energy of amino acids side chain analogues. Next, we systematically evaluated the accuracy and efficiency of different potential functions-CHARMM36, CHARMM36m and the scaled CHARMM36 in predicting the weak self-association of ubiquitin and the formation of a heterodimer between EphA2 SAM domain and SHIP2 SAM domain. Finally, we examine the protein association pathways of the 25 cases where native complexes of E-SAM: S-SAM are formed and find that a multi-stage process is at play.

## METHODS

### Estimation of Solvation Free Energy

The standard free energy perturbation method was used to calculate the solvation free energy of amino acid side chain analogues:^43–46^

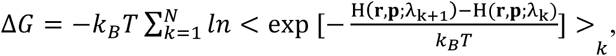

where *λ* is a scale factor (and a small perturbation) that gradually tune the potential energy function of a system. Specifically, in our simulation, the potential energy function between the side chain analogue and water was first tuned gradually from λ = 1 to 0 by a decrement δλ = 0.02 (forward simulation) in a 2 ns simulation; and was gradually turned back on from λ = 0 to 1 in an additional simulation period of 2.0 ns (backward simulation). 50 intermediate λ-states or “windows” (N) are produced with an interval of δλ = 0.02. In order to avoid charge singularities at λ = 0 or 1, a soft-core potential was used to smoothly switch off the interactions. Specifically, outgoing atoms see their electrostatic interactions with the environment decoupled during λ = 0 to 0.5, while van der Waals interactions are gradually decoupled from λ = 0 to 1. Therefore, the electrostatic interaction is fully switched off before the van der Waals interaction is tuned to 0. Each window was simulated for 40 ps, with a 4 ps long equilibration and a 36 ps simulation for data collection. Overall, for each calculation of the solvation free energy for a specific amino acid analogue, the system was first equilibrated for 2.0 ns. A subsequent 4.0 ns of the simulation (forward and backward simulation) was used to perform the free energy perturbation. Three repeats were performed to estimate the uncertainties. The solvation free energy of individual amino acid side chain analogues was calculated with different CHARMM potential functions.^47^ In particular, we carried out calculations with the original CHARMM36 potential function,^48^ and a scaled potential function CHARMM36s (see below), and made comparisons with the results of Bert et al^44^ performed with the CHARMM22 potential function and with experimentally measured values.^49^ It should be noted that CHARMM36m made changes on the CMAP correction terms^50^ to optimize the protein main chain fold. In addition, the interaction between guanidinium (-NH3 group of ARG) and of the carboxylate (-COO group of ASP/GLU) was reduced by modifying vdw radii from 0.3550 nm to 0.3637 nm.^36^ None of the two changes affect the solvation of an individual amino acid. Therefore, the solvation free energy of individual amino acid side chain analogues are the same between CHARMM36 and CHARMM36m.

### Set-up of Simulation System of Protein Association

Next, we evaluated the accuracy and efficiency in predicting protein-protein interactions with three different sets of potential functions-CHARMM36,^48^ CHARMM36m^36^ and CHARMM36s. Two model systems were tested: the self-association of ubiquitin and the hetero-association of E-SAM and S-SAM. For ubiquitin system, two ubiquitins [pdb id. 1ubq] were separated by 40 Å at the beginning of simulation. Following an equilibrium for 20 ns, we performed 2 μs production simulations on the Anton2 supercomputer for each of the potential functions. For E-SAM: S-SAM association with a moderately strong binding affinity, instead of performing a long production simulation, we carried out multiple independent simulations (60 simulations each, for three sets of parameters) to enhance the computational sampling. The E-SAM and S-SAM domains of the NMR derived complex [pdb.id 2kso, ref. 51] were also separated to a distance of 40 Å between their center of mass. Both SAM domains were then rotated randomly with respect to their center of mass in each of the 60 simulations to produce different initial orientations for each simulation with CHARMM36. In simulations with CHARMM36s and CHARMM36m, we used the same set-ups of initial orientations between two SAMs as in simulations with CHARMM36, thus there are no bias between three sets of parameters in terms of starting simulation configurations. The nearest distance between any atoms of the two SAMs are about 6-18 Å, mostly (in 48 of 60 simulations) larger than 10 Å. The whole system (ubiquitin or E-SAM: S-SAM) was solvated by TIP3P water^52^ in a simulation box of 90×90×90 Å^3^, setting up periodic boundary conditions. Sodium and Chloride ions were added to a near-physiological concentration of 150 mM and to neutralize the system. Side chain groups were charged as expected at pH 7.0 (Histidine with hydrogen on delta carbon-HSD/HID, was used). In total, the simulation box contained about 71,000 atoms. The system set-up followed previous protocols (e.g. initial heating and equilibration for 40 ps and 1 ns, respectively [e.g. 22]). Each simulation was 250-300 ns long.

### Simulation Parameter and Analysis

In all simulations, the van der Waals (vdW) potential was truncated at 12 Å and smoothly shifted to zero between 10 and 12 Å to reduce cutoff noise. The Particle-Mesh Ewald (PME) method was used for calculating the long distance electrostatic interactions.^53^ The SHAKE algorithm was applied for all covalent bonds to hydrogen. The time step was set as 2 fs. The systems were coupled to a thermostat at 300 K and a barostat at 1 bar with the Langevin scheme.^54^ All the systems were simulated using the NAMD/2.12 package.^55^ The simulations of ubiquitin were further transferred to Anton supercomputer^56^ to simulate the self-association at a longer time scale of 2 μs. Analysis (interface-RMSD, residues-residues contact, surface electrostatics, orientational angle, pair interactions, and solvent accessible surface) were done with VMD or in NAMD with in-lab scripts (for descriptions see figure legends/main text). The first time of contact between the two SAM domains was noted when any atoms of the two structures came within 4 Å from one another. The same cutoff was used to construct contact maps. In calculation of interface-RMSD (i-RMSD), the Cα position of interface residues 949-962 of E-SAM and residues 1215-1239 of S-SAM were used for an otherwise standard RMSD calculation, with respect to the average Cα location in the NMR ensemble.^51^ A protein complex with an i-RMSD < 4.0 Å with respect to native complex is considered as native-like, given that this value encompasses the different orientation states of the two SAM domains relative to one another from the original solution NMR analysis.^51^ Incidentally, 4 Å is also the cut-off chosen for i-RMSD to indicate acceptable protein complex structures in the Critical Assessment of Predicted Interactions (CAPRI) experiment.^24^ The similarities of residue-residue contact maps were made between current simulations of protein association and prior simulations starting from a native complexes of E-SAM: S-SAM. Native contacts were considered as interactions between: L951: H1219/N1220/G1221, P952: N1220, G953: N1220/G1221/W1222, H954: G1221, K956: W1222/D1235, R957: D1224/F1227, Y960: D1230 (numbers x:y refers to residue x in EphA2- or E-SAM, y to SHIP2- or S-SAM). The similarities were calculated by comparing these interactions in simulations with different potential functions to the prevalence of these interactions in simulations starting from a native complex (results from ref. 22). The unscreened vdW and electrostatic interactions for Fig. 7 and Fig. S4a-d were calculated with a cutoff of 32 Å in absence of solvent and ions.

## RESULTS

### Optimization of Solvation Free Energy of Amino Acid Side Chains by Scaling Protein Atom-Water Interactions

Side chains of amino acids are dominant groups that mediate the protein-protein interactions. However, in many of the prevalent potential functions (e.g. CHARMM and AMBER), the solvation free energy of amino acid side chain analogue is underestimated relative to the experimental value. The differences vary from −0.2 kcal/mol to as large as −2.5 kcal/mol. The inaccuracy is particularly large for amino acids that typically constitute “hot spots” at protein-protein interface (PPI) such as hydrophobic residue ILE and aromatic residues TYR and TPR (Fig. 1). The underestimated solvation promotes protein-protein association. A scaling of side chain: water interaction is a straightforward and empirical way to increase the solvation of amino acid side chain. We made modification on the CHARMM36 potential function parameter by scaling the solute-solvent vdW interaction potential energy. Firstly, the interactions between -CH2, –CH3 group and water is increased by a small scale factor of 1.03 (i.e. an increase of 3%). These changes, with the scale factor optimized manually in these and the subsequent side chains, help to better fit the solvation of hydrophobic amino acids of ALA, VAL, LEU, ILE. As –CH2, and – CH3 groups are the only scalable groups in these amino acid side chains, they are scaled first, forming the basis as they occur in other side chains. To further increase the solvation of polar residue, a scale factor of 1.10 was applied to polar S, N and C at the terminal of SER, THR, MET, CYS. The solvation free energy was not calculated for charged residues where solvation free energy calculations are made difficult by the frequent presence of counter ions.^43–46^ However, due to stereo-similarity at the side chain level to their ASN/GLN polar counterparts, the same scaling was also applied to basic and acidic residues LYS, ARG, ASP and GLU. At last, a scale factor of 1.05 was applied to the benzene group in PHE, TYR and TRP. It improves the solvation of benzene; in the meantime, it increases the solvation of the aromatic amino acids. All the changes applied to individual amino acids are shown in Figure S1.

**Figure 1:**
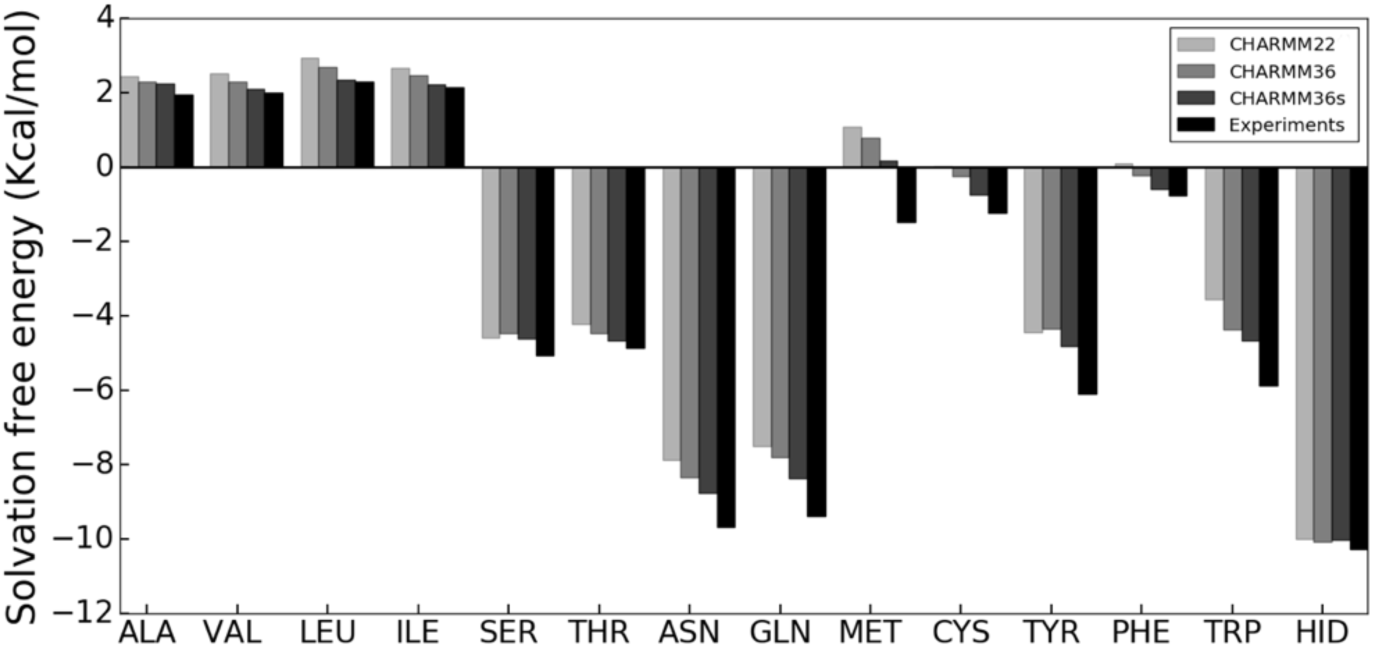
Solvation Free Energy of Amino Acid Side Chain Analogues. Comparisons are made with experimental data (ref. 49) and the molecular dynamics simulation results with CHARMM22 (by Shirts et al., ref. 44), CHARMM36, and CHARMM36s. HID indicates Histidine with H atom bonded to N^δ^.

As shown in Fig.1, all of the amino acids achieve better solvation with vdW scaling of the solute-water interactions in comparison to CHARMM22 (results from ref. 44) and CHARMM36/36m. On average, the solvation (excluding ALA and protonated His, HSD) is increased by 0.36 kcal/mol for all residues with CHARMM36s (see also Table S1). In two pioneering studies by Best et al and Feig et al, similar scaling on solute-water interactions has been applied.^14,37^ However, in these studies a much higher vdW scaling factor of 1.10 was applied to all protein atoms. The solvation free energy calculation showed that a 1.10 scaling affords too much solvation for the amino acids, already apparent in the study of Best et al.^37^ In addition, modifications on the main chain atoms likely made the protein fold unstable (data not shown). Therefore, in this study, the scale was only applied to some of the amino acid side chain atoms; and overall, a much lower scaling factor (mostly 1.03, and few atoms with 1.05 and 1.10) was used. For brevity, we denoted the modified CHARMM36 with scaled solute-water interactions as CHARMM36s.

### Simulations of weak Non-Specific Interactions between Ubiquitin

Based on the NMR results, ubiquitin has a non-covalent self-association with a very weak binding affinity, K_d_ around 4.8 mM,^57^ which is likely to manifest in frequent dissociation and association events which should be captured even in a short simulation. However, the study by Abriata et al. showed that in molecular dynamics simulation ubiquitin has a rather strong self-association, aggregates easily and in a non-physiological manner.^32^ Similarly, in our simulation, with the original CHARMM36 potential function ubiquitin associates into a dimer shortly after the start of the simulation, and does not dissociate over the remainder of the 2 µs simulation (Fig. 2a). With improved solvation of the amino acid side chains in CHARMM36s, “overly sticky” protein-protein interactions are partially released (Fig. 2b). The protein complex undergoes a few dissociation-(re)association events, but eventually forms a relatively stable complex from 1.5 to 2 µs. With CHARMM36m, the protein complex undergoes rather frequent dissociation-(re)association events (Fig. 2c). Fig. 2d plots the distribution of distance between the centers of mass of proteins. This distance is within 3 nm for 90.5%, 77.6% and 34.2% of time for two ubiquitins in the simulation trajectories, using CHARMM36, CHARMM36s and CHARMM36m respectively (by comparison, the radius of gyration of ubiquitin is 1.2 nm). Based on the experimental K_d_ value above, the concentration of ubiquitin (4.8 mM) in the simulation, and supposing that the kinetics of association and dissociation are fast, the fraction of ubiquitin dimer should be 50%. Clearly, both potential function parameter changes improve the predictions of the monomer-dimer equilibrium of ubiquitin, but CHARMM36s still overestimates the bound population whereas CHARMM36m may slightly underestimate it. The NMR study indicated that residues 4-12, 42-51, and 62-71 are the major contacts at the protein-protein interface (due to the high flexibility, res. 72 to 76 were not analyzed^57^). In the simulations with CHARMM36, the prevalence of PPIs at res. 4-12 or 62-71 is not observed. Simulations with CHAMM36s and CHARMM36m both show some interactions at res. 4-12 and 62-71, but CHAMMR36s shows only a very low frequency for these interactions. Surprisingly, compared to CHARMM36/-s, CHARMM36m shows rather wide-spread/non-specific interactions across the entire protein.

**Figure 2:**
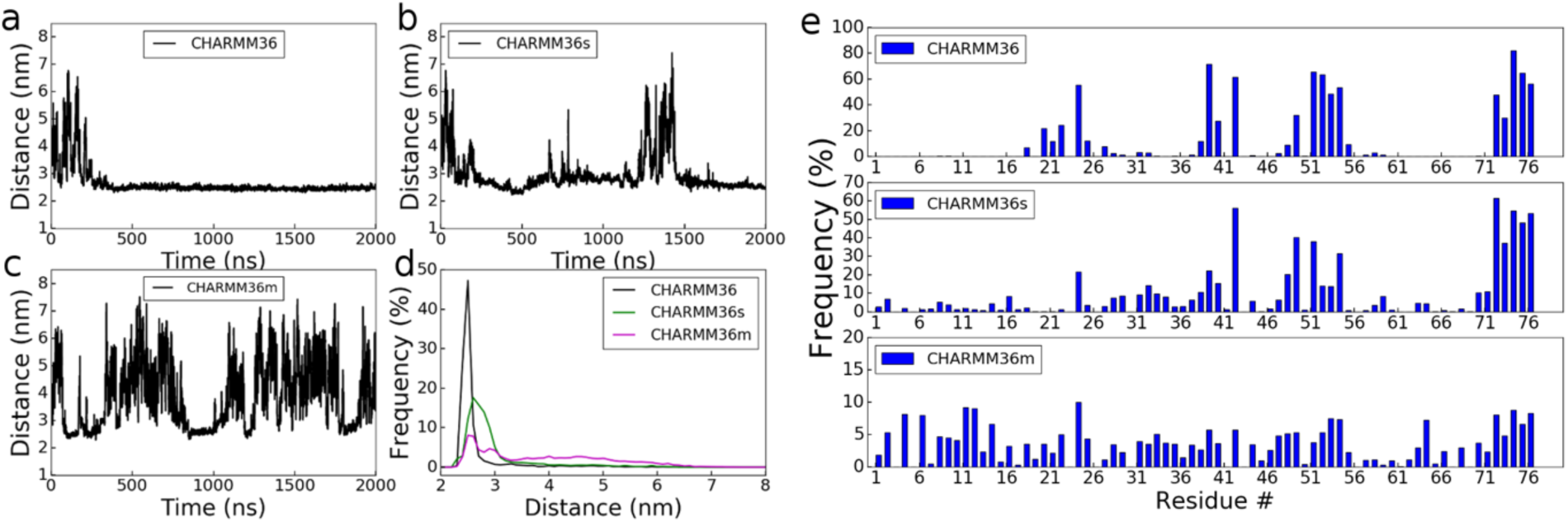
Self-association of ubiquitin over time course of a 2,000 ns simulation, run with different potential functions. (a-c) Time evolution of distance between the centers of mass of two ubiquitins. (d) Distribution of distance between the centers of mass of the two ubiquitins. (e) Frequency of individual residues at the ubiquitin-ubiquitin interface.

There is only one parameter correction in CHARMM36m that is directly related to protein-protein interactions: the vdW minimum potential between atom type of NC2 (ARG) and OC (GLU/ASP) is shifted from 0.3550 nm to 0.3637 nm. The influence of this correction is dramatic. While the change is small for the vdW potential energy, the change in inter-atom distance is large in terms of electrostatics, considering that HC (connected to NC2) and OC have partial charges of 0.46 and −0.76 e respectively. The longer the equilibrium distance between HC and OC, the greater is the decrease in the electrostatic interactions between the ARG and GLU/ASP side chains. A rough estimation (without considering the vicinity of water or possible counter ions) indicates that the interactions between the –COO and the – C(NH_2_)_2_ group is increased by 3.24 kcal/mol from −107.48 kcal/mol with CHARMM36 to −104.24 kcal/mol with CHARMM36m (Figure S2). In the self-association of ubiquitin, the association involves guandinium-carboxylate interactions as well as interactions between LYS and ASP/GLU. Therefore, the reduction of the guanidinium-carboxylate interaction in CHARMM36m largely attenuates its self-association. However, considering the fraction of ubiquitin dimers is less than that expected in simulations with CHARMM36m, the distribution of residues at PPI is rather even and the reduction of guanidinium-carboxylate interactions may overly weaken protein associations. However, this point still needs to be cautiously examined in the other systems and further examined via free energy calculations of protein-protein binding.

### Overall Enhanced Prediction of E-SAM: S-SAM Native Protein Complexes with Optimized Potential Function Parameters

Due to our previous extensive experimental and computational studies of EphA2-SAM: SHIP2 SAM complex, we chose this complex, with moderate binding affinity with a K_d_ of 2 µM, to examine a protein-protein association process in detail. Using the original potential function, CHARMM36, we found that even long simulations could not dislodge several of the trapped states that exist shortly after association (data not shown). Raising the simulation temperature (from 300K to 323K or 350K) or increasing the salt concentration (from 150 mM to 1.5 M NaCl) did not overcome this problem either. In order to increase simulation sampling, we performed 60 simulations each of 250-300 ns for each set of simulation parameters, CHARMM36, /-s and /-m.

Fig. 3 plots the distribution of the interface RMSD (i-RMSD) defined by the root mean square distance to native contacts (see Methods). Clearly, the simulations with modified potential function parameters have a better performance in sampling native-like structures than simulations with the original CHARMM36 potential function. Specifically, with original CHARMM36, only 4 of 60 simulations yields a native complex. However, with the modified potential functions in CHARMM36s, the simulations yield 10 native complexes; with CHARMM36m, 11 native complexes were observed. So the yield of native structures was increased from 6.7% to 16.7%, and 18.3% respectively. A significant difference between the CHARMM36s and /-m potentials is that the former has several structures in the range of 5-6 Å i-RMSD whereas the latter has very few, suggesting that, for CHARMM36m, the free energy minimum of the native complex is well separated by having energy barriers that restrict access to surrounding minima.

**Figure 3:**
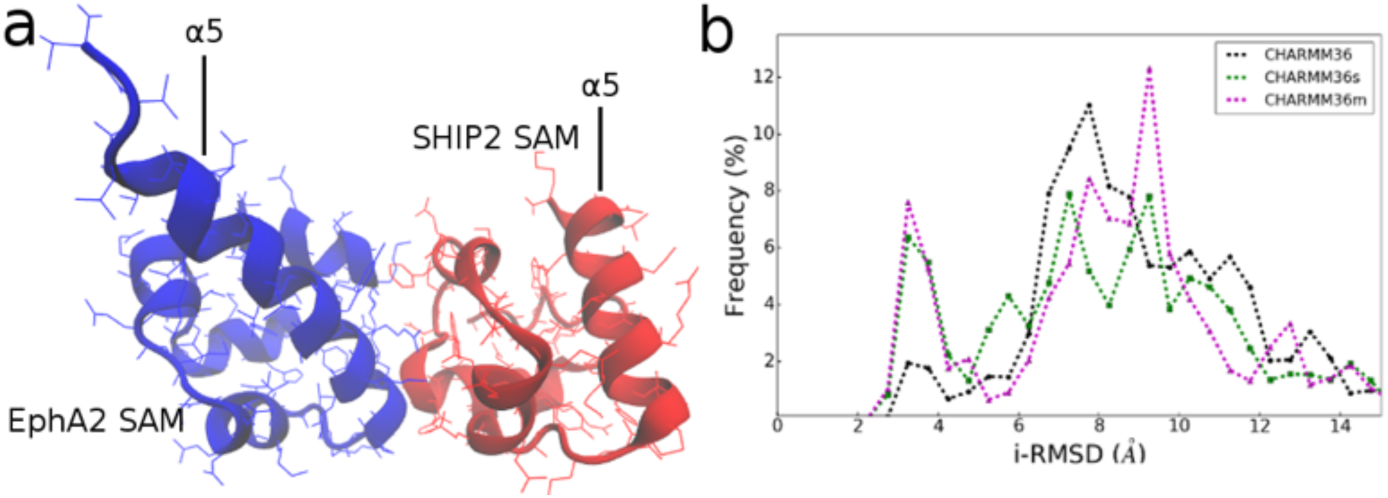
Enhanced prediction of native complex of E-SAM: S-SAM. (a) Heterodimer of E-SAM: S-SAM solved by NMR.^51^ E-SAM is in blue color and S-SAM is in red color throughout the paper. (b) Distribution of i-RMSD for different simulation sets (i-RMSD for structures in the last 50 ns of the 60 simulations).

Figure 4 compares the difference in residue-residue contact patterns (for the last 50 ns of the simulations), between simulations with original and modified parameters. Counting the major interacting residues in the native complex, the contact similarity between the native configuration and the latter part of the association simulations with CHARMM36, /-s and /-m are 5.5%, 12.5% and 12.5% respectively (see Methods for calculation details). As non-specific/transient contacts are averaged, the pattern of native contacts (see M&M) is more apparent in the simulations with modified parameters. Furthermore, remarkably, the major region of native contacts, E-SAM res. 952-957: S-SAM res. 1220-1224, 1227 and 1235 is particularly enhanced in simulations with the modified potential function. Even in cases where a native-like complex is not formed, one of the two proteins typically uses its correct native-like/high affinity interface for making contacts. The enhanced sampling with the improved potential functions indicates that the inaccuracy of CHARMM36 potential function accounts at least in part for the low yield rate of native structure in original simulations.

**Figure 4:**
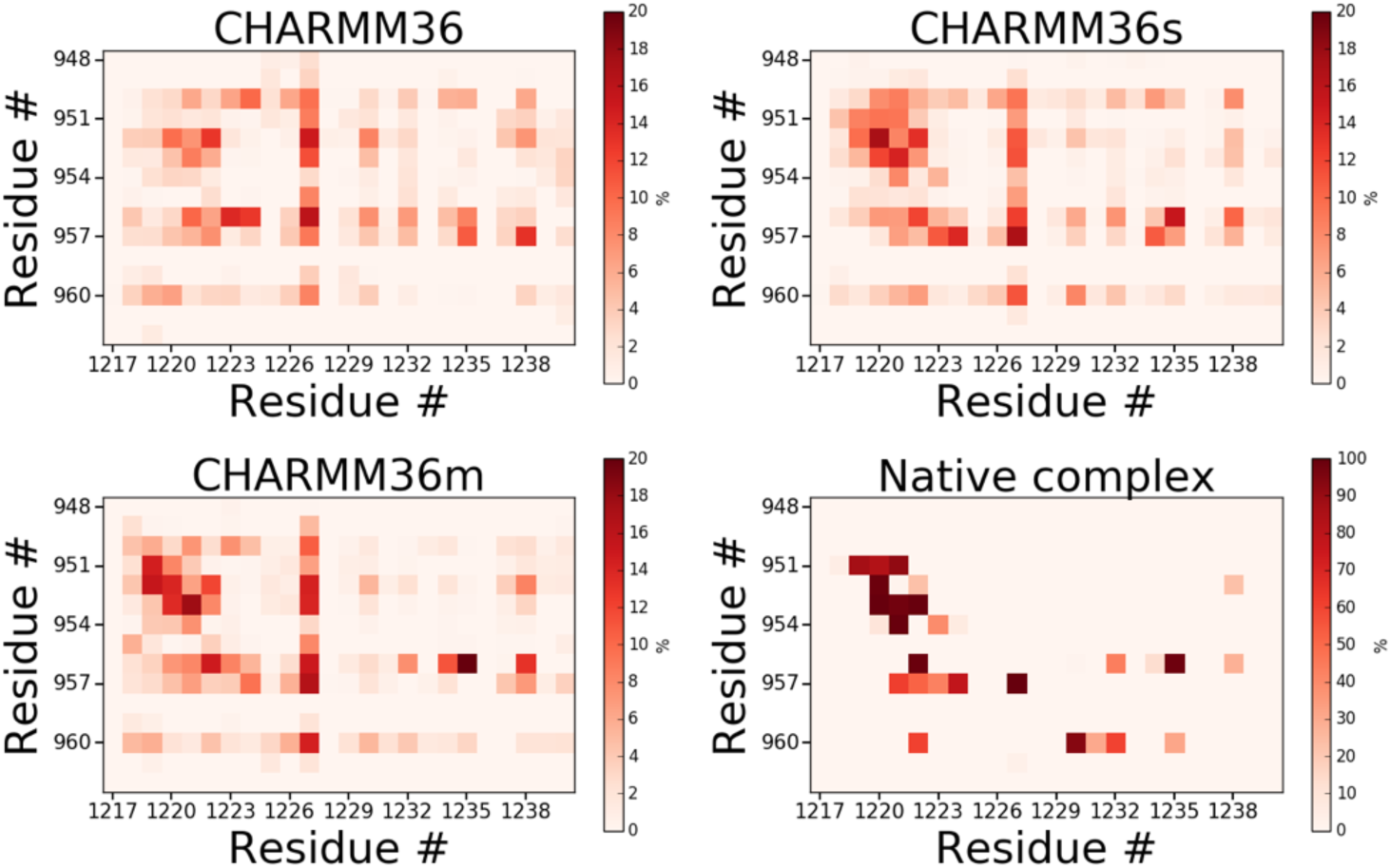
Residue-residue contact maps for different simulation sets (subset of map showing regions of native-like interactions only). The last 50 ns of the simulation trajectories were used. Contact map at the bottom right) was obtained based on the simulation trajectories starting from native complex (ref. 22). A much smaller scale bar is used in for simulations with CHARMM36, /-s, /-m as many of the simulation do not form native complexes, so the maximum of occupancy of native contact are typically less than 20%. This contrasts simulations which are started from a native complex which have some contacts near 100% occupancy on this timescale.^22^ Outside of this region the non-native contacts are spread very widely and are of low frequency.

### Comparing the Sensitivity of the Early Stages of Association to Changes in the Potential Function

Given that the potential function modifications overall weaken the protein-protein interactions, we may anticipate a delay in the time to first substantial contact. Indeed, simulations with CHARMM36s and /-m provide a good number of trajectories where contacts are made later than in simulations run with the original potential function (Fig. 5a). However, the simulations with CHARMM36s and /-m also have significantly more dissociation events, 12 and 15 respectively, than simulations with CHARMM36 (8 dissociation events). In each simulation set, there is only one native-like complex that is formed after a protein-dissociation event, suggesting that by itself a dissociation does not create conditions for a favorable subsequent association process.

**Figure 5:**
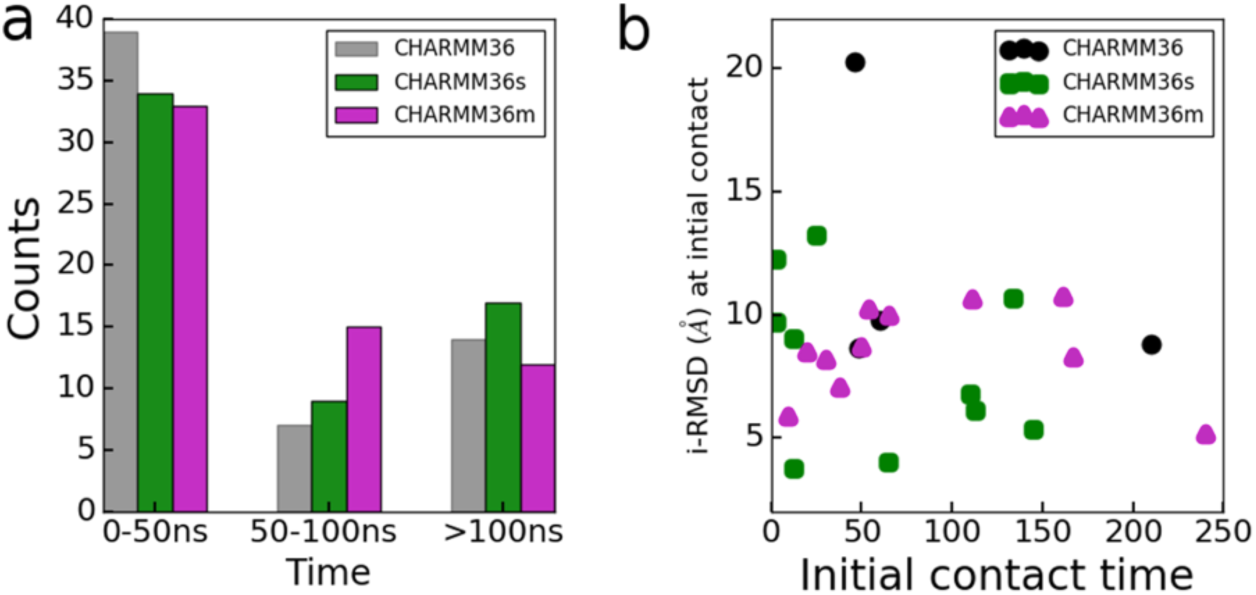
Difference between simulations with original and modified potential function. (a) Initial protein-protein contact time for 60 independent simulations. (b) i-RMSD value at the initial contact time (shown for simulations that get to a native complex).

For simulations that lead to a native complex, we plot the i-RMSD at the initial contact time (Fig. 5b). Again, a cut-off of 4 Å for i-RMSD is set as the criteria of the formation of a native protein complex. In order to assess the closeness of a bound complex at the initial contact time to the native complex, we further distinguished two different situations depending whether the deviation of the i-RMSD from the 4 Å criteria is within 5 Å of or not (i-RMSD value less than or larger than 9Å). It shows that at the initial contact time, 2, 5 and 5 of the complexes have already positioned themselves into a configuration that is in vicinity of the native complex in the CHARMM36, -/s and /-m simulations respectively. Such a better pre-orientation in the simulations with the modified potential may be observed because during the approach of the two proteins, they are less easily trapped in non-native configurations, but also they have more time to pre-orientate. On the other hand, once the proteins are associated, with the modified potential function, the proteins could also more easily undergo orientation changes to form a native complex. In the other cases (2, 5, 6 for CHARMM36, -/s and /-m respectively), at the time of the initial contact, the bound complex is not close to the native configuration (big i-RMSD value). In order to form a native complex, the bound complex needs to adjust themselves to get to the native configuration.

The initial contact between E-SAM and S-SAM are made within 50 ns in more than half of the simulations, irrespective of parameter modifications (Fig. 5a). The preferential distribution of the centers of mass of S-SAM relative to E-SAM that results from such pre-orientation and translation events is shown in Fig. 6a for simulations with CHARMM36. The S-SAM move towards to E-SAM, and by 50 ns mostly concentrate on one region made up of helices α4 and α5 (and less so α1) of E-SAM (the coordinate frames are superimposed on E-SAM in all molecular snapshots shown). Intriguingly, while initial contacts are confined to residues ∼945-960 on the E-SAM side, no such dramatic confinement exists on the side of S-SAM (Fig. 6b). The pre-orientation of S-SAM relative to E-SAM is expected to be directed by electrostatic interactions. As shown in Fig. 6c, the primary binding region of S-SAM has many negative charges and thus a negative surface electrostatic potential (esp. far right). However, the region of E-SAM which contains helices α2 and α3 (residues 922-938) is also negatively charged. Therefore, the interaction between these regions is unfavorable, reflected by the absence of center of mass localization of S-SAM around this region of E-SAM. Instead, the E-SAM region containing helix α4 and α5 is positively charged (electrostatic surface depicted on top left). The two proteins associate with each other through this preferred interface which shows a strong electrostatic complementarity. This supports the viewpoint that an electrostatic pre-orientation (or electrostatic steering) of the domains is involved to direct the early stages of protein association. Negative charged side chains are more extensively distributed on the S-SAM surface, even though negative charge is more concentrated on one side of the protein. This may explain why there is no dramatic confinement of E-SAM relative to the S-SAM surface. These patterns of pre-orientation, over the first 50 ns, only change slightly upon modification of the potential function; results for CHARMM36m and /-s are shown in the SI (Fig. S3). This result is not too surprising as the modifications involve only short range vdW contacts directly (and in the case of /-s only vdW contacts to water). Thus, the effect of the potential function change is more pronounced at shorter interaction distances, i.e. once the proteins have associated. Another important point is, however, that the distributions of center of mass of the SAM domains around each other is very similar over the first 10 ns and is extensive, showing a wide sampling of starting orientations of the 60 simulations in each condition.

**Figure 6:**
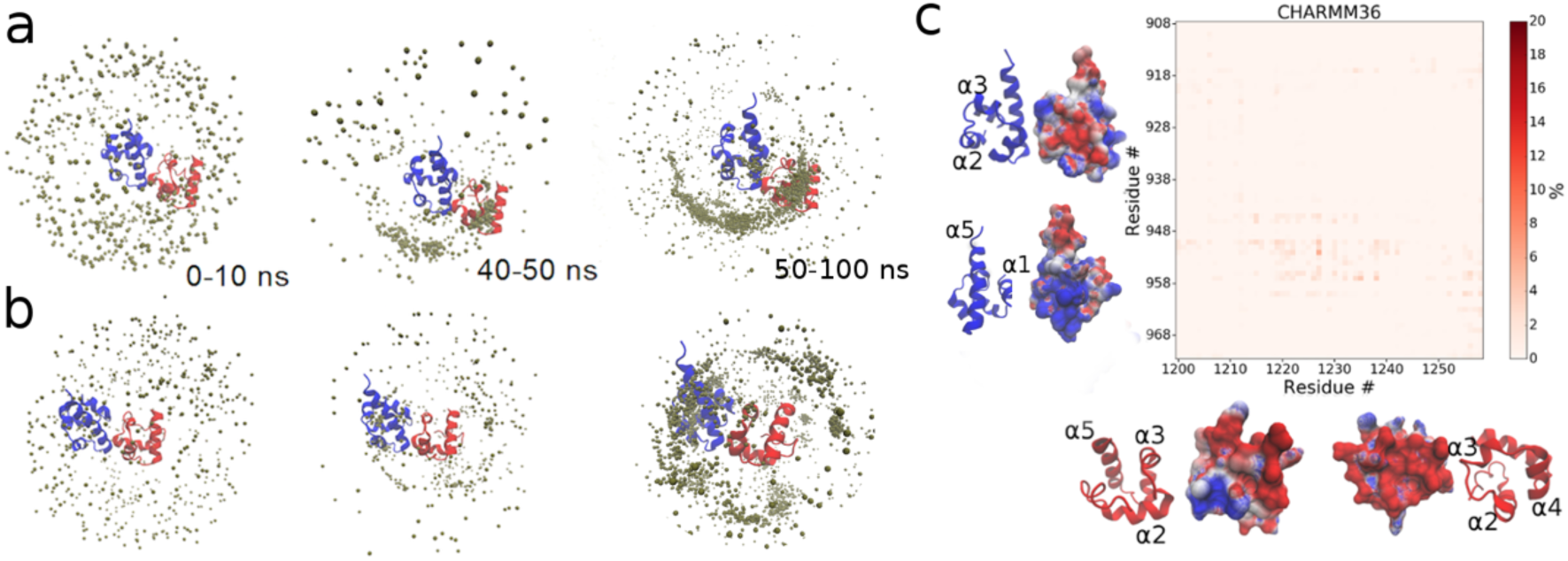
Pre-orientation during the early stage of protein association. (a-b) The early stages of E-SAM and S-SAM association for simulation with the original potential function. Pre-orientation of S-SAM (E-SAM) relative to E-SAM (S-SAM). Proteins are superimposed on E-SAM (a) or S-SAM (b), respectively. The center of mass of S-SAM (or E-SAM) is denoted by small beads. (c) E-SAM: S-SAM residue-residue contact map during the early stage of protein-protein association in simulations with original potential function. Simulation trajectories of 0-50 ns are used to construct the contact maps, averaged over 60 trajectories. The color scale (=contact occupancy; max near ∼4.8%) is from grey to red as indicated. The contact occupancy is still low, as not all proteins are extensively interacting with the same residues. Electrostatic surfaces are shown alongside the respective E- and S-SAM residue axes, with SAM orientations, beside them as main chain cartoons. The electrostatic surface was calculated with the APBS module of VMD and is shown at ± 4 kT from negative (red), neutral (white) to positive electrostatic potential (blue). Results from the other two potential functions are shown as as Figure S3 in SI.

### Transition Pathways in the Associated Proteins

Fig. 7 shows time evolution of a representative protein-protein association (simulation #17 with CHARMM36). At around 140 ns, two proteins first associate. As the protein get closer, more residue-residue contacts are established (Fig. 7a), the value of i-RMSD decreases (Fig. 7c), more solvent available surface area is buried (Fig. 7d), and the pair interaction gets more favorable (Fig. 7e), but still it does not represent a native complex. The initial contact is maintained for about 50 ns. At a time around 190 ns, the protein dissociated from the non-native structure and established contacts again at 215 ns. As shown in Fig. 7b, after making protein-protein contacts at 215 ns, S-SAM slides along the surface of E-SAM (215-233 ns) and eventually adjusts itself to a native structure through a major rotation (233-249 ns). Along with the protein sliding and rotation, the i-RMSD gradually decreases below the 4 Å criteria (Fig. 7c). In the meantime, a more buried surface accessible surface area (SASA) and lower potential energy for electrostatic pair interactions are achieved (Fig. 7d-e). The low value for vdW interactions is consistent with our prior simulations for this system. We suggested elsewhere that the association of E-SAM and S-SAM is less hydrophobic (only one side chain is fully desolvated) and many side chains are still partially solvated.^25^ Protein-protein electrostatic interactions drive the orientation of the proteins to form the complex, but the actual complex formation involves a sizeable desolvation of the protein-protein interface region as a major driving force, reversing the process we noted upon SAM domain dissociation.^25^ Some residues at the SAM domain interfaces (in this transition, E-SAM G953/R957 and S-SAM H1219) could be critical for the transitions, acting as pivots, consistent with the concept of “anchoring residue” introduced by Camaco and Colleagues.^58^

**Figure 7:**
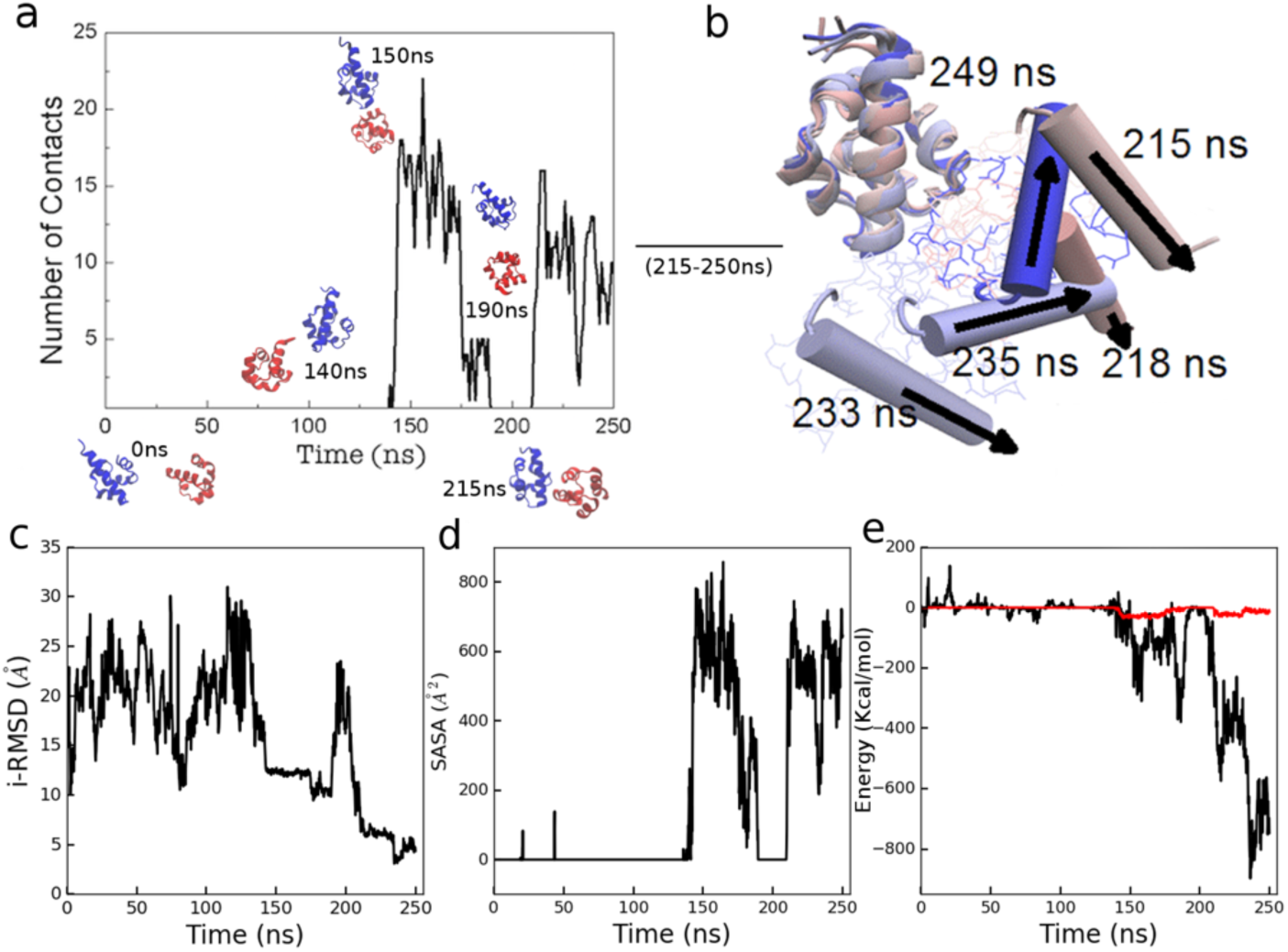
A protein association process (from simulation #17 with CHARMM36). (a) Number of contacts between residues of the E-SAM and S-SAM domain. Molecular images show representative snapshots. (b) A dynamic transition towards to the native structure occurs in the bound state. Helix 5 of S-SAM are shown in cartoon representation. (c) i-RMSD and (d) buried surface accessible surface area (SASA) of the E-SAM and S-SAM. (e) Unscreened protein-protein Electrostatic (black) and vdW interactions (red) between E-SAM and S-SAM. (Further examples, corresponding to the 4 simulations in Fig. 8 are plotted in Fig. S4a-d).

Figure 8a shows other examples of the protein-protein association process (with CHARMM36s and /-m), in which large orientation changes are needed to transform to a native complex. Similarly, S-SAM needs to slide on the surface of E-SAM but it also involves self-rotations to adjust to the native complex especially for sim. #15 with CHARMM36m. We measured the angular displacement (*ϕ*) of the center of mass of the S-SAM domain (relative to the orientation of E-SAM) between the initial contact time and the finial simulation time by making a superposition of both coordinate frames on the E-SAM structure. Thus angle ϕ describes the surface movement of S-SAM relative to E-SAM, but S-SAM can also self-rotate while it moves relative to E-SAM. To obtain information about rotations of S-SAM relative to itself, in the same way, we measured the angular displacement (φ) of E-SAM (relative to S-SAM) between the initial contact time and final simulation time, when the structures are superimposed on S-SAM. For both simulations shown in Fig. 8a, the orientation change indicated by ϕ and φ are larger than 45°. Fig. 8b shows two other representative simulations performed with CHARMM36s and CHARMM36m. In these two cases, there are less obvious orientational adjustments to reach a native complex. At the initial contact, which occurs around 145 ns and 9 ns, respectively, the i-RMSD (5.3 Å and 5.9 Å respectively) is already very close to a native complex. The relative orientation change is correspondingly small (less than 30°). Therefore, in these two cases, the formation of a native complex is the result of a good pre-orientation. Fig. 8c shows change of relative orientation of two SAM domains between initial contact and final structure for simulations that form a native complex. In 0, 5 and 6 of the simulations with CHARMM36, /-s and /-m, respectively, both angles (ϕ and φ) have changes less than 45°, which indicates relatively small orientation adjustments. In 2, 5, 5 simulations with CHARMM36, /-s and /-m. respectively, the orientation change of either ϕ or φ is larger than 45° but less than 90° (see also Table S2 in Supporting Information). Besides, in 2 simulations with CHARMM36, ϕ is larger than 90° for unknown reason. Taken together, the modification on the potential function partially eliminates overly sticky between E-SAM: S-SAM, by allowing a better pre-orientation and easier surface movement, at least over short reorientations of one domain relative to the other. Both processes provide mechanisms to release complexes from trapped states, leading to an enhanced prediction of native-like structures.

**Figure 8:**
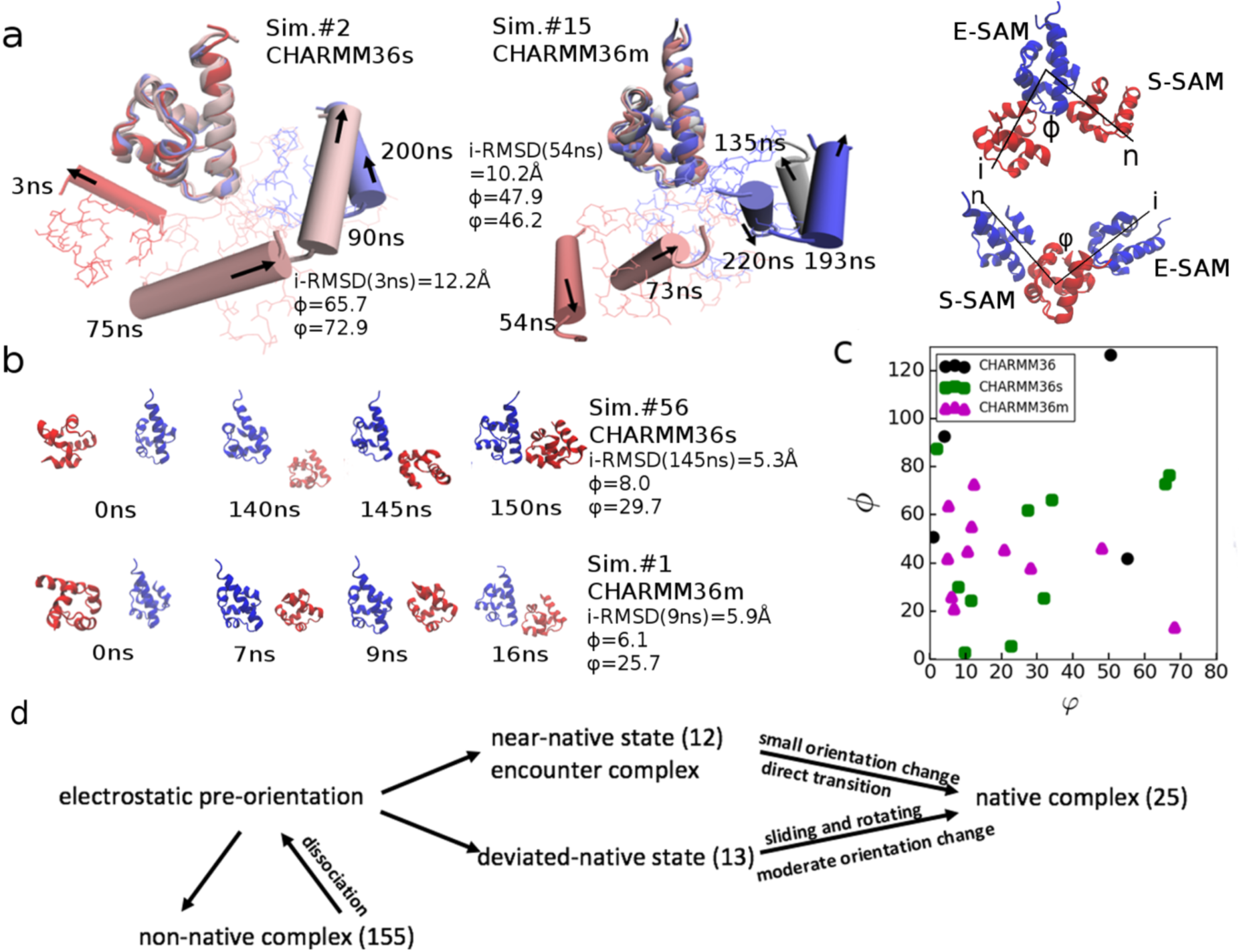
Protein association process between E-SAM and S-SAM. (a) Snapshots show a dynamic transition towards to the native structure that occurs in the bound state. Helix 5 of S-SAM is shown in cartoon, representing (in part) the orientation of the domain. One example is from Simulation #2, run with CHARMM36s, and the other from Simulation #15 run with CHARMM36m. The right top molecular diagram show the definition of change of relative orientation of two SAM domains between initial contact and final structure. ϕ denotes the cross angle between the two vectors---the one connects the center of mass of E-SAM to the center of mass of S-SAM at initial contact time, and the other one connects the center of mass of E-SAM to the center of mass of S-SAM at the final simulation time. ϕ indicates the orientation changes of S-SAM relative to E-SAM (E-SAM is superimposed), Similarly, φ denotes the orientation change of E-SAM relative to S-SAM when S-SAM is superimposed over the trajectories. i indicates the initial structure while the n indicates the final native structure. (b) Snapshot for a pre-orientation process of the unbound SAM domains. Shown for Simulation #56, run with CHARMM36s and simulation #1, run with CHARMM36m. (c) The change of relative orientation for simulations that get to a native complex. See Table S2 for results of individual simulations. (d) Summary of pathways of all 25 protein-protein associations leading to a native complex.

Despite of using improved potential functions, 155 of the 180 simulations result in non-native complex (Fig. 8d). 35 dissociation events were recorded, although only 3 of them lead to a native complex via rebinding. If the initial association is too distant from the native structure (for example, S-SAM binding to the opposite side of E-SAM), in principle, the proteins may also directly move around one another by rotational translation/sliding along the binding partner’s protein surface to get to the native structure. This relative movement of the two proteins involves a considerable amount of making and breaking of intermediary/new residue-residue interactions. So a trapped sub-stable complex may better convert to a native-like configuration by going through a dissociation-(re)association process, rather than by an extensive movement along the other domains surface to which it is already bound. While in the current simulations, only 3 native complex were formed via a dissociation-rebinding process, frequent dissociation-rebinding likely increase the fraction of native complex eventually over longer periods of simulation time. In most of the 25 simulations (in which a native complex was formed) the S-SAM is typically already positioned in vicinity of interfaces made of α4, α5 and α1 of E-SAM at the initial contact; so overall the complex is not very distant from the native structure. Based on the analysis of i-RMSD, in 12 simulations, the i-RMSD at the initial contact time is less than 5Å away from the 4 Å criteria of forming a native complex, while in the another 13 simulations, the deviation of i-RMSD at the initial contact time is larger than 9Å. Based on the analysis of orientational angle change, in 11 simulations, both orientation angles (∅ and *φ*) experience changes less than 45° between the first contact and final structures. In another 14 simulations, at least one angle (∅ and *φ*) changes more than 45° but typically less than 90°. We distinguished two different protein associations as shown in Fig. 8d. In the first case, since the protein configuration of the complex already position very close to the native one, only small orientation adjustment is needed to transform into a native state (examples shown in Fig. 8b). The concept of productive “encounter complex” indicates protein complex that are in vicinity of native state, which could easily transform into native state. In this sense, these well pre-orientated protein complexes resemble the concept of a likely productive encounter complex. However, it should be noted that even when the complex has a pre-orientation that is close to native complex, with i-RMSD <9Å at the initial contact time, the complex can still become trapped in a non-native structure over the remainder of the simulation time. In the second case, the protein complex is relatively far (but still not too distant) from a native complex at the beginning. Thus, in order to form a native complex, moderate orientation adjustments involving protein sliding and rotating with each other are needed to reach the native state (examples shown in Fig. 7b and Fig. 8a).

## DISCUSSION

This study shows that the formation of native protein-protein interactions can be enhanced by apparently subtle changes to the potential function. CHARMM36m is clearly beneficial for the predictions of the native SAM-SAM complex. By reducing the guanidinium-carboxylate interactions between ARG and ASP/GLU, the electrostatic attraction between two proteins is reduced and this allows for a greater number of dissociations. But our work suggests that the reduction of guanidinium-carboxylate interaction may go further than necessary. Intramolecular interactions via ARG:ASP/GLU within a singular protein should also weaken and this may affect the stability of a protein. The recent benchmark study by D. E. Research of multiple popular potential functions indicated that some proteins show reduced stability with CHARMM36m.^36^ However, CHARMM36 and CHARMM36m underestimate the solvation free energy of amino acid side chains. Quantum chemistry calculations, a main driver in potential function parameterization, may not fully appreciate the roles of water and thus result in an underestimated solvation of amino acids. It should be noted that improvements in the water model used, did not completely improve the solvation of amino acid side chains. Except for TIP3P_MOD (which is rarely used and is subjected to a large change of the vdW radius and strength of water), water models TIP3P, TIP4P and SPC all showed underestimated solvation of amino acids.^45^ A scaling of solute-water interactions is an empirical way to eliminate the bias for solute-solute and water-water interactions. We used a set of small scaling factors and only applied the scaling to the interactions between amino acids side chains and water. By scaling solute-water vdW interactions (in CHARMM36s) we bring the solvation free energies of amino acid side chains closer to experimental values. Importantly, we also found that the increased protein solvation is beneficial for the formation of native PPIs. However, the new parameters have been evaluated here for only two examples of protein association. While they significantly improve the prediction of ubiquitin self- and SAM domain hetero dimer-association, the proteins are small and have only small hydrophobic surface patches. It is, therefore unclear how the modification of the potential function may affect protein-protein association in the other systems and how it may affect the stability of a wide variety of individual proteins. We expect that the increased solvation likely affect processes where the solvation is of key importance, for example, the conformations of intrinsically disordered proteins (IDPs), or protein-protein association which are largely mediated by hydrophobic or aromatic residues. As interactions with water are scaled up the potential function will also likely reduce the partitioning of amino acids into membrane and thus influence protein-membrane interactions. These effects need to be examined in future studies to evaluate the influence of solvation optimization on different processes.

The still relatively modest yield of native structures is possibly due to inaccuracies that remain in other aspects of the current potential function parameters, but it is also likely that much longer sampling on the order of tens, if not hundreds of microseconds is required for finding native-like protein complexes.^16,59^ Ideally, in protein-protein complex formation, the native structure corresponds to a deep but narrow free energy well, while the non-native sub-stable structures are more wildly spread and have broader minima that are (again ideally) connected to the native well. For example, we notice that the distribution of i-RMSD over the last 50 ns of CHARMMM36m trajectories (Fig. 4) have a minimum between approx. 4.5-6 Å, suggesting that complexes in this range are at the crest of a funnel that quickly moves them either towards or away from the native-like complex. Beyond this region the energy landscape is likely to be an even more complicated terrain. This complexity, likely makes the process of finding and even maintaining the native complex structure especially hard in simulations that start from separated proteins and only use physical parameters. Indeed, a number of laboratories are pursuing the use of knowledge-based potentials or combinations of knowledge-based and physical potentials.^60–62^ Enhanced sampling methods, such as temperature- or Hamiltonian-based replica exchange or simulated annealing molecular dynamics have been used in more extensive searches of configurational space of individual proteins or peptides,^16,59^ but are just beginning to be used in the prediction of protein-protein association.^63,64^

Our study suggests that CHARMM36m and, alternatively, a CHAMM36s with scaled solute-solvent interactions, partially remove current inaccuracies in the potential function. By plotting the pair interaction versus the i-RMSD (see Fig. S5a), we can clearly see a positive correlation between low potential energy states and near-native complex for i-RMSD > 10Å in all simulations. Other energy functions which include solvation free energy and maybe even dynamics/entropy (free energy functions) would be more appropriate, but also more difficult to calculate. These kinds of analysis could be used for predicting the native complex without prior knowledge about the protein-protein interface. The native complex could be inferred based on the overall popularity as well as low-energy properties of a specific cluster of configurational states. It is intriguing however, that if we just plot the data for the 25 successful trajectories, we see a funnel with a clear minimum at around 3A, especially in the case of CHARMM36s (the original potential function has a broad but rather flat minimum, while CHARMM36m has a broad but slanted minimum, again showing a correlation with i-RMSD). This observation suggests that the simulations which go the native complex, do so in a well defined energy funnel. However, the accuracy of the prediction of PPIs still depend on probability of transitioning into this funnel (as opposed to becoming trapped in local energy minima), which is reflected by the overall yield rate of native complex. If the yield rate could reach to, for instance, 30%-50%, by clustering the protein complex and examining energetics information, it is likely to give a reliable prediction of native protein complex. The current brute force all-atom simulations show that an improvement in the potential function itself (without enhanced sampling) already can significantly increase the accuracy of prediction of protein-protein association. And as discussed above, there is likely further scope for additional improvements in the potential function, but also in sampling (enhanced sampling or much longer simulations). With a further optimized potential function and increased computational power, a precise prediction of native complex via molecular dynamics simulation should be expected. *{see notes added below}

Previously we reported extensive simulations of the SAM-SAM heterodimer in two papers. One paper showed that in several simulations one SAM domain underwent configurational reorientations relative to the other, while keeping the same residues at the protein-protein interface, just changing the identity of the interacting charged residues.^22^ However, in the SI of that paper we also showed several simulations which did not undergo such transitions but became stuck, with one interface slightly moved away from the other. We now think that at least some of these mismatches developed because we used the CHARMM36 force field and the parameters were too sticky. However, a second paper on the dissociation process of two SAM complexes with salt bridge pair swap mutants,^25^ revealed some of the features (pivoting and rolling in place of one protein relative to the other) seen here in the association simulations.

The formation of native complexes largely occurs through a pre-orientation process that could position the S-SAM in vicinity of the PPI of native complex. Are the association and dissociation processes of the SAM-SAM complex the reverse of one another? We could show that the protein-protein association indeed involves less sliding or reorientation if the encounter complex is “productive”, that is, it is already close to a native complex. Therefore, in these few cases, the association is not the reverse as the dissociation process we observed previously, where (albeit with affinity reduced mutants) transitions to a more weakly binding surface appeared to be obligatory before the proteins could separate.^25^ Generally, and despite modification of the potential function, the mechanism of binding and finding the native state complex is overall similar to the reverse of the dissociation trajectories we had with complex destabilizing mutants.^25^ Similar to the viewing an association in reverse, coming out of a deep energy minimum for dissociation seems best accomplished by a surface sliding or pivoting movement to a weaker binding interface first. We should caution, however, that our conclusion may not be general as it is based on the study of a SAM-SAM complex, which has highly complementary electrostatics and yet –as a cell signaling/adaptor domain interaction-has a much weaker binding affinity than classical protein complexes (e.g. Barnase-Barstar which interact very tightly with a deep, funnel-like energy landscape^65^). The greater number of dissociation-(re-)association events could in principle also help the formation of a better pre-orientated complex but this is more rarely observed. It is intriguing that the effects of quite different parameter changes are similar in the ease of dissociation and of configurational changes within the complex, while most of the successful simulations go to the native state without a dissociation step. This suggests that the rate determining interactions rely on similar close range contacts. These observations are in agreement with the findings of Feig, Sugita and colleagues who, in all-atom simulations of the cytoplasm of a bacterial cell discovered that metabolites spend a considerable time rolling on protein surfaces, rather than diffusing between proteins.^66^

Since the early pioneering work of R. Wade and colleagues, it is known that the formation of encounter states can be fast and is well modeled by Brownian Dynamics which employ rigid particles with an emphasis on long and medium range interactions.^8–10^ On one hand, a close correspondence between theoretical and experimental parameters, such as salt dependence and electrostatics modifying mutants is possible (most experiments measure the formation of the encounter state, rather than of the native state). On the other hand, only a very small fraction of encounter complexes has a sufficient number of native-like contacts, to give small i-RMSD values and convert to native complex structures. In many circumstances, the second step, that of a configurational transition of the already bound state, is thought to be rate limiting for the formation of the native complex. Recently, spin-labeling and other types of NMR experiments have opened the door to a residue specific characterization of the initial bound encounter, if not transient, intermediate non-native bound states of protein complexex.^67,68^ Such experiments have started on the SAM: SAM complex in our laboratory.

## CONCLUSION and PERSPECTIVE

In summary, we investigated the protein-protein association with an original potential function and two slightly modified potential functions, utilizing classical all atom simulations. With the original CHARMM36 potential function, the native-like structures of the protein complex are obtained in 6.7% of the all-atom unbiased molecular dynamics simulations of the EphA2 (E-)SAM: SHIP2 (S-)SAM heterodimer within 250 ns. By improving the solvation free energy of amino acids side chains in a potential function we denote CHARMM36s, the prediction of native complex is increased to 16.7%. Due to a decrease of electrostatic attraction between ARG and GLU/ASP, the recently published CHAMMR36m potential function significantly reduces the protein-protein interactions between ubiquitin homodimers, and also of the E-SAM: S-SAM complex, yielding native complexes in 18.3% of simulations. In all trajectories, electrostatic interactions between charged residues are a key force in directing the initial protein-protein association. In order to form a native complex quickly, one protein ideally pre-orientates to a position that is in vicinity of the native protein-protein interface of the other. From such a pre-orientated positon, the initial complex could in some cases directly transform to the native complex with small orientation changes; Alternatively, in other cases, the formation of a native complex involves moderate changes. Overall, the study provides a rich picture on mechanisms of protein-protein complex formation in a small model system. In future, computational modeling by itself should be able to predict protein-protein interactions with high accuracy. This will help to direct the site mutagenesis experiments, and accelerate the modeling of structure-level protein interactomes. The reliable prediction of protein complexes would also greatly reduce the experimental expense (incl. number of experiments), especially considering that only a minor fraction of structures is currently available for protein interactomes (predicted to be of at least 130,000 protein complexes for humans^69^). With increased computing power, physical protein association trajectories may go to 100s of microseconds (if not several milliseconds) and in combination with further optimized potential functions should yield a higher rate of native complex formation.

## *ADDED NOTE

After submission of an earlier version of this manuscript on bioRxiv, two other groups submitted all-atom simulations on protein association, both using enhanced sampling. Specifically, in a study by D. E. SHAW research, the researchers developed an enhanced sampling method that could very accurately predict native complexes in lengthy MD simulations. The simulations use a strategy that -by constantly scaling down the protein-protein interactions (both vdW and electrostatic potential function parameters)-help the protein escape from non-native complexes; however, the extent of parameter modification is likely protein dependent (http://www.biorxiv.org/content/10.1101/303370v1). A study by the Chong lab. uses a weighted ensemble method to guide the protein association along a pathway ultimately guided by a reduction of i-RMSD (https://www.biorxiv.org/content/10.1101/453985v1).

## ACKNOWLEDGEMENTS

This work was supported by NIGMS grant R01GM112491 to the Buck lab. and used the Extreme Science and Engineering Discovery Environment (XSEDE) Stampede at Texas Advanced Computing Center (TACC), Ohio Supercomputer Center (OSC) as well as local computing resource in the core facility for Advanced Research Computing at Case Western Reserve University. Anton Computer time was provided by the Pittsburgh Supercomputing Center (PSC) through Grant R01GM116961 from the National Institutes of Health.

## ASSOCIATED CONTENT

The Supporting Information is available free of charge on the ACS Publications website at http://pubs.acs.org.

Graph of atoms with scaled solvent-solute interaction (Fig. S1); list of solvation free energy of amino acid side chain analogue (Table S1); difference in guanidium interaction between CHARMM36 and CHARMM36m (Fig. S2); list of i-RMSD, *ϕ* and *φ* for all simulations (Table S2); residue-residue-residue contact map (Fig. S3); time evolution of residue-residue contact, i-RMSD, buried SASA, pair interactions shown for representative simulations with CHARMM36m and 36s (Fig. S4a-d); and correlation between pairwise protein-protein potential energy and i-RMSD (Fig. S5a,b).

The authors declare no competing financial interests.

---

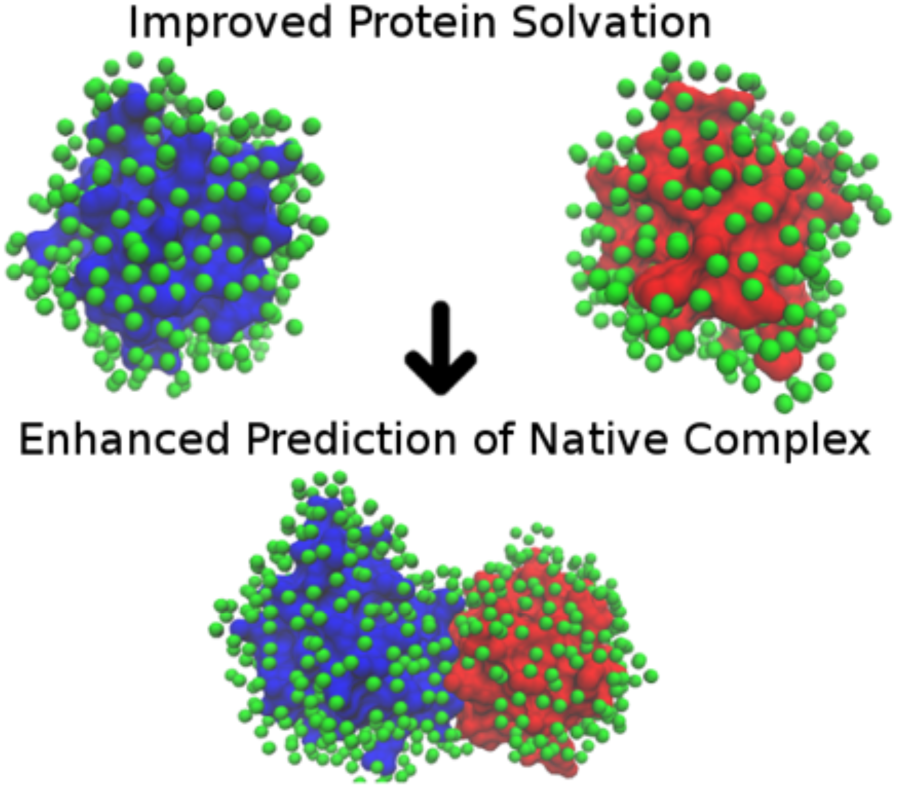
For Table of Contents Only

