## Supplemental Information for "Modified Potential Functions Result in Enhanced Predictions of a Protein Complex by All-Atom MD Simulations, Confirming a Step-wise Association Process for Native PPIs"

Zhen-lu Li[1] and Matthias Buck[1,2*]

[1]Department of Physiology and Biophysics, Case Western Reserve University, School of Medicine, 10900 Euclid Avenue, Cleveland, Ohio 44106, U. S. A.

[2]Departments of Pharmacology, of Neurosciences and Case Comprehensive Cancer Center, Case Western Reserve University, School of Medicine, 10900 Euclid Avenue, Cleveland, Ohio 44106, U. S. A.

*

Figure S1: Atoms affected by scaling of solute-solvent interactions for the CHARMM36s potential function.

(1) CHARMM atom type: **CT2 CT2A HA2 CT3 HA3** scaled by 1.03

```
ALA
   |
HN-N
   |    HB1
   |    /
HA-CA-CB-HB2
   |    \
   |    HB3
 O=C
   |
```

```
LEU
   |       HD11 HD12
HN-N          | /
   |   HB1   CD1--HD13
   |   |    /
HA-CA--CB--CG-HG
   |   |    \
   |   HB2   CD2--HD23
 O=C          | \
   |       HD21 HD22
```

```
ILE
   |    HG21 HG
HN-N      | /
   | CG2--HG23
   |    /
HA-CA--CB-HB    HD1
   |    \        /
   |     CG1--CD--HD2
 O=C    / \      \
   | HG11 HG12  HD3
```

```
VAL
   |   HG11 HG12
HN-N     | /
   |    CG1--HG13
   |    /
HA-CA—CB
   |    \
   |    CG2--HG21
 O=C    /  \
   |  HG11  HG12
```

(2) CHARMM atom type: CC, OH1, NH2, NC2 scaled by 1.10

```
SER (OH1)
   |
HN-N
   |   HB1
   |   |
HA-CA—CB--OG
   |   |     \
   |   HB2    HG1
 O=C
   |
```

```
GLN (CC, NH2)
   |
HN-N
   |  HB1 HG1 OE1  HE21
   |   |   |   ||   /
HA-CA--CB--CG--CD--NE2
   |   |   |         \
   |  HB2 HG2        HE22
 O=C
   |
```

```
CYS (S)
   |
HN-N
   |    HB1
   |    |
HA-CA--CB--SG
   |    |     \
   |    HB2    HG1
 O=C
   |
```

```
THR (OH1)
   |
HN-N
   |    OG1-HG1
   |   /
HA-CA--CB-HB
   |   \
   |    CG2--HG21
 O=C     / \
   |   HG21 HG22
```

```
ASN(CC, NH2)
   |
HN-N
   |  HB1 OD1   HD21
   |   |   ||   /
HA-CA--CB--CG--ND2
   |   |        \
   |  HB2       HD22
 O=C
   |
```

```
MET (S)
   |
HN-N
   |  HB1 HG1     HE1
   |   |   |       |
HA-CA--CB--CG--SD--CE--HE3
   |   |   |       |
   |  HB2 HG2     HE2
 O=C
   |
```

```
ARG (NC2)
   |                     HH11
HN-N                      |
   |  HB1 HG1 HD1 HE     NH1-HH12
   |   |   |   |   |    //
HA-CA--CB--CG--CD—NE= CZ
   |   |   |   |        \(+)
   |  HB2 HG2 HD2        NH2-HH22
 O=C                      |
   |                     HH21
```

```
LYS (NH3)
   |
HN-N
   |  HB1 HG1 HD1 HE1     HZ1
   |   |   |   |   |      /
HA-CA--CB--CG--CD--CE--NZ--HZ2 (+)
   |   |   |   |   |      \
   |  HB2 HG2 HD2 HE2     HZ3
 O=C
   |
```

```
ASP (CC)
   |
HN-N
   |   HB1   OD1
   |   |    //
HA-CA— CB--CG
   |   |    \
   |   HB2   OD2(-)
 O=C
   |
```

```
GLU (CC)
   |
HN-N
   |  HB1 HG1   OE1
   |   |   |   //
HA-CA--CB--CG--CD
   |   |   |   \
   |  HB2 HG2   OE2(-)
 O=C
   |
```

(3) CHARMM atom type: **CA, CPT, CAI** scaled by 1.05

```
HSD / HID
   |          HD1     HE1
HN-N           |      /
   |   HB1     ND1--CE1
   |   |      /       ||
HA-CA--CB--CG         ||
   |   |      \\      ||
   |   HB2     CD2--NE2
 O=C          |
   |          HD2
```

```
PHE(CA)
   |        HD1  HE1
HN-N         |    |
   |   HB1  CD1--CE1
   |   |    //     \\
HA-CA--CB--CG      CZ--HZ
   |   |    \  __  /
   |   HB2  CD2--CE2
 O=C         |    |
   |        HD2  HE2
```

```
TRP (CA,CPT,CAI)
   |                  HE3
HN-N                   |
   |   HB1            CE3
   |   |             /   \\
HA-CA--CB---CG-----CD2   CZ3-HZ3
   |   |    ||     ||     |
   |   HB2  CD1    CE2   CH2-HH2
 O=C       /   \   / \   //
   |     HD1     NE1   CZ2
                  |     |
                 HE1   HZ2
```

```
TYR (CA)
   |        HD1  HE1
HN-N         |    |
   |   HB1  CD1--CE1
   |   |   //      \\
HA-CA--CB--CG      CZ--OH
   |   |    \  __  /     \
   |   HB2  CD2--CE2     HH
 O=C         |    |
   |        HD2  HE2
```

*Table S1: Solvation Free Energy of Amino Acid Side Chain Analogue (Kcal/mol).*

| Amino Acid Side Chain Analogue | Experiment (ref. 50) | CHARMM36 | CAHRMM36s | CHARMM22 (ref. 45) |
|---|---|---|---|---|
| ALA (methane) | 1.94 | 2.29 ± 0.07 | 2.24 ± 0.03 | 2.44 |
| VAL (propane) | 1.99 | 2.28 ± 0.04 | 2.08 ± 0.10 | 2.52 |
| LEU (isobutene) | 2.28 | 2.68 ± 0.05 | 2.33 ± 0.06 | 2.94 |
| ILE (butane) | 2.15 | 2.46 ± 0.09 | 2.21 ± 0.12 | 2.67 |
| SER (methanol) | -5.06 | -4.48 ± 0.11 | -4.63 ± 0.08 | -4.59 |
| THR (ethanol) | -4.88 | -4.47 ± 0.08 | -4.68 ± 0.09 | -4.22 |
| ASN (methanethiol) | -9.68 | -8.38 ± 0.09 | -8.77 ± 0.12 | -7.89 |
| GLN (propionamide) | -9.38 | -7.82 ± 0.07 | -8.39 ± 0.01 | -7.51 |
| MET (methyle ethyl sulfide ) | -1.48 | 0.78 ± 0.07 | 0.18 ± 0.12 | 1.08 |
| CYS (methanethiol) | -1.24 | -0.25 ± 0.03 | -0.74 ± 0.02 | 0.02 |
| Benzene | -0.87 | -0.34 ± 0.05 | -0.72 ± 0.11 | NA |
| TYR (p-cresol) | -6.11 | -4.35 ± 0.21 | -4.83 ± 0.19 | -4.46 |
| PHE (toluene) | -0.76 | -0.22 ± 0.08 | -0.60 ± 0.20 | 0.09 |
| TRP (methylindole) | -5.88 | -4.38 ± 0.08 | -4.66 ± 0.14 | -3.57 |
| HID (methylimidazole) | -10.27 | -10.08 ± 0.05 | -10.04 ± 0.06 | -10.00 |

Figure S2: Estimation of unscreened guanidium interaction consisting of Lennart-Jones (LJ) and Coulomb (Coul) interactions with the potential function parameters of (a) CHARMM36 and (b) CHARMM36m. The LJ and Coul interactions are calculated along with the distance between the terminal Carbon atom of a GLU residue and an ARG residue (c). The position of each atom is based on the equilibrium bond length and bond angel. (d) Comparison of the total interaction energy of guanidium from a) and b) i.e. between CHARMM36 and CHARMM36m.

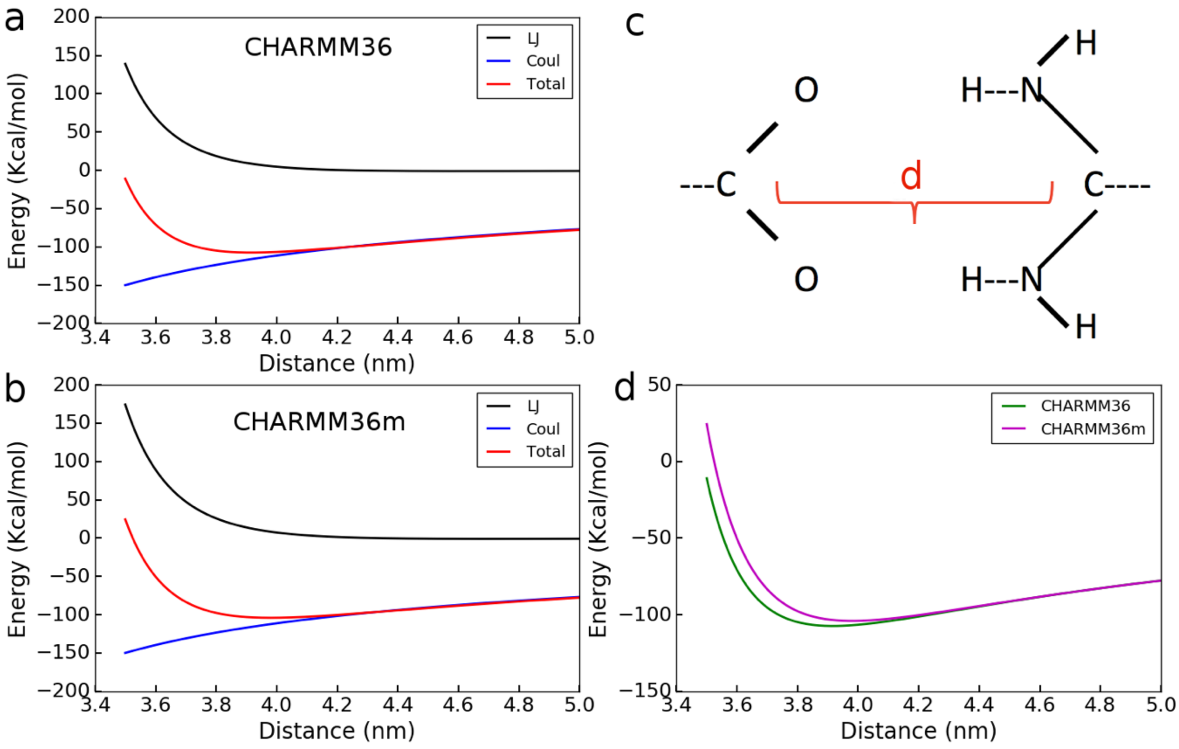

Table S2: List of initial contact time, i-RMSD (relative to native complex) at initial contact and final simulation time, and orientational change (ϕ and φ) between initial time point of association and final simulation frame. Rows with values in **bold**, indicate simulations that form a native complex.

| Simu. # | Time (ns) | i-RMSD$_i$ (Å) | i-RMSD$_f$ (Å) | $\phi$ (°) | $\varphi$ (°) |
|---|---|---|---|---|---|
| CHARMM36 | | | | | |
| 1 | 11.0 | 7.5 | 7.8 | 16.4 | 30.6 |
| 2 | 45.0 | 13.6 | 6.7 | 24.3 | 70.7 |
| 3 | 4.0 | 10.9 | 10.7 | 17.4 | 11.1 |
| 4 | 29.0 | 21.5 | 13.2 | 9.5 | 9.5 |
| 5 | 68.0 | 15.4 | 15.1 | 32.0 | 11.5 |
| 6 | 7.0 | 12.8 | 7.8 | 39.4 | 53.5 |
| 7 | 104.0 | 6.3 | 11.6 | 21.0 | 125.2 |
| 8 | 12.0 | 8.1 | 7.1 | 17.3 | 15.5 |
| 9 | 7.0 | 10.4 | 8.0 | 23.6 | 17.4 |
| 10 | 4.0 | 11.8 | 10.0 | 26.9 | 12.4 |
| **11** | **46.0** | **20.3** | **3.8** | **50.4** | **126.9** |
| 12 | 116.0 | 11.7 | 13.2 | 21.6 | 44.6 |
| 13 | 185.0 | 9.2 | 12.9 | 17.3 | 60.4 |
| 14 | 11.0 | 17.0 | 15.5 | 10.7 | 103.3 |
| **15** | **48.0** | **8.6** | **3.7** | **4.0** | **92.9** |
| 16 | 36.0 | 6.9 | 8.5 | 31.1 | 46.4 |
| **17** | **210.0** | **8.8** | **4.4** | **0.9** | **50.9** |
| 18 | 3.0 | 10.9 | 10.1 | 11.5 | 18.4 |
| 19 | 141.0 | 13.6 | 7.9 | 16.7 | 13.6 |
| 20 | 117.0 | 20.2 | 6.0 | 118.9 | 75.4 |
| 21 | 17.0 | 8.7 | 6.8 | 43.3 | 7.3 |
| 22 | 69.0 | 16.7 | 14.6 | 11.0 | 3.1 |
| 23 | 14.0 | 9.1 | 8.7 | 0.1 | 9.8 |
| 24 | 36.0 | 11.1 | 8.9 | 22.5 | 120.0 |
| 25 | 47.0 | 15.3 | 11.1 | 23.5 | 35.2 |
| 26 | 94.0 | 8.7 | 4.7 | 6.1 | 59.4 |
| 27 | 111.0 | 15.3 | 13.2 | 12.4 | 25.8 |
| 28 | 18.0 | 7.7 | 8.2 | 15.3 | 2.0 |
| 29 | 148.0 | 13.9 | 7.5 | 86.2 | 58.0 |
| 30 | 113.0 | 11.6 | 9.1 | 27.3 | 10.0 |
| 31 | 66.0 | 9.4 | 6.8 | 50.3 | 22.1 |
| 32 | 163.0 | 9.4 | 7.2 | 7.8 | 27.3 |
| **33** | **60.0** | **9.8** | **3.4** | **55.2** | **41.9** |
| 34 | 14.0 | 8.4 | 7.4 | 25.2 | 16.4 |
| 35 | 115.0 | 10.4 | 8.6 | 30.3 | 1.5 |
| 36 | 27.0 | 8.1 | 7.1 | 16.8 | 45.8 |
| 37 | 25.0 | 14.7 | 9.8 | 49.0 | 30.8 |
| 38 | 16.0 | 9.7 | 7.6 | 2.5 | 5.3 |
| 39 | 11.0 | 10.0 | 9.1 | 20.8 | 39.6 |
| 40 | 9.0 | 11.8 | 10.5 | 70.1 | 47.4 |
| 41 | 105.0 | 9.6 | 8.1 | 5.1 | 22.5 |
| 42 | 11.0 | 11.9 | 12.0 | 30.0 | 37.3 |
| 43 | 29.0 | 14.7 | 6.5 | 38.5 | 120.4 |
| 44 | 23.0 | 10.5 | 10.7 | 29.4 | 24.6 |
| 45 | 11.0 | 10.9 | 11.7 | 9.5 | 53.0 |

| | | | | | |
|---|---|---|---|---|---|
| 46 | 206.0 | 16.8 | 10.3 | 56.3 | 102.3 |
| 47 | 77.0 | 11.0 | 7.5 | 32.4 | 43.9 |
| 48 | 213.0 | 13.1 | 7.5 | 44.5 | 30.0 |
| 49 | 4.0 | 16.2 | 7.3 | 144.6 | 39.6 |
| 50 | 28.0 | 12.8 | 13.1 | 5.6 | 40.9 |
| 51 | 3.0 | 8.1 | 10.7 | 34.6 | 26.6 |
| 52 | 16.0 | 21.4 | 6.0 | 59.1 | 133.5 |
| 53 | 5.0 | 14.4 | 11.3 | 17.2 | 55.3 |
| 54 | 24.0 | 10.8 | 11.2 | 2.8 | 18.4 |
| 55 | 74.0 | 11.1 | 7.5 | 8.1 | 37.7 |
| 56 | 6.0 | 18.3 | 7.2 | 53.5 | 123.2 |
| 57 | 35.0 | 11.8 | 7.9 | 52.6 | 40.6 |
| 58 | 4.0 | 9.4 | 7.1 | 9.5 | 65.0 |
| 59 | 22.0 | 16.3 | 7.6 | 87.1 | 65.2 |
| 60 | 8.0 | 15.0 | 12.1 | 32.5 | 117.9 |
| CHARMM36s | | | | | |
| 1 | 69.0 | 10.6 | 4.7 | 2.1 | 26.7 |
| **2** | **3.0** | **12.2** | **3.0** | **65.7** | **72.9** |
| **3** | **65.0** | **4.0** | **4.9** | **22.9** | **5.1** |
| 4 | 9.0 | 17.8 | 16.9 | 4.0 | 23.6 |
| 5 | 156.0 | 18.5 | 17.3 | 2.8 | 6.3 |
| **6** | **13.0** | **3.7** | **4.0** | **9.7** | **2.6** |
| 7 | 105.0 | 8.6 | 7.6 | 17.7 | 17.3 |
| 8 | 230.0 | 17.8 | 10.8 | 44.9 | 38.9 |
| 9 | 4.0 | 11.8 | 6.6 | 3.3 | 32.9 |
| 10 | 183.0 | 11.7 | 7.6 | 12.9 | 112.0 |
| **11** | **110.0** | **6.7** | **4.7** | **31.9** | **25.3** |
| 12 | 3.0 | 19.9 | 10.9 | 5.0 | 46.8 |
| 13 | 159.0 | 12.2 | 12.1 | 23.8 | 0.6 |
| 14 | 115.0 | 8.9 | 9.5 | 54.1 | 28.0 |
| 15 | 118.0 | 20.9 | 9.3 | 47.2 | 49.8 |
| 16 | 23.0 | 9.7 | 5.6 | 13.1 | 57.9 |
| 17 | 14.0 | 15.0 | 14.1 | 10.0 | 14.8 |
| 18 | 18.0 | 14.2 | 8.7 | 13.0 | 103.1 |
| **19** | **113.0** | **6.1** | **4.2** | **11.6** | **24.3** |
| 20 | 66.0 | 11.7 | 7.1 | 22.2 | 22.0 |
| 21 | 8.0 | 17.9 | 6.1 | 51.3 | 88.3 |
| 22 | 44.0 | 9.2 | 9.1 | 21.2 | 11.0 |
| 23 | 36.0 | 12.2 | 9.2 | 33.8 | 66.0 |
| 24 | 11.0 | 14.7 | 5.9 | 1.0 | 86.9 |
| 25 | 72.0 | 16.7 | 9.3 | 49.9 | 2.5 |
| 26 | 15.0 | 11.0 | 8.7 | 20.3 | 39.3 |
| 27 | 75.0 | 11.7 | 7.2 | 4.7 | 14.7 |
| 28 | 6.0 | 8.0 | 10.4 | 18.8 | 37.1 |
| 29 | 9.0 | 11.5 | 10.6 | 79.6 | 34.2 |
| **30** | **13.0** | **9.0** | **3.6** | **34.2** | **66.3** |
| 31 | 8.0 | 13.8 | 10.8 | 2.8 | 22.2 |
| 32 | 138.0 | 14.9 | 11.1 | 3.5 | 36.9 |
| 33 | 12.0 | 8.9 | 9.2 | 2.9 | 4.4 |
| 34 | 250.0 | 11.5 | 9.1 | 25.7 | 37.0 |
| **35** | **25.0** | **13.2** | **3.2** | **2.0** | **87.5** |

| | | | | | |
|---|---|---|---|---|---|
| 36 | 4.0 | 11.1 | 7.1 | 5.5 | 18.7 |
| 37 | 25.0 | 14.6 | 6.1 | 37.2 | 67.0 |
| 38 | 38.0 | 7.7 | 7.3 | 17.1 | 0.5 |
| 39 | 60.0 | 9.6 | 7.5 | 13.1 | 49.7 |
| **40** | **134.0** | **10.6** | **3.4** | **27.5** | **61.9** |
| 41 | 37.0 | 14.3 | 8.0 | 64.3 | 6.7 |
| 42 | 86.0 | 14.1 | 14.0 | 32.5 | 17.0 |
| 43 | 4.0 | 21.1 | 14.4 | 2.2 | 10.3 |
| 44 | 103.0 | 9.3 | 9.8 | 31.8 | 8.1 |
| 45 | 28.0 | 14.9 | 16.4 | 35.8 | 61.5 |
| 46 | 185.0 | 16.3 | 10.9 | 54.5 | 87.3 |
| 47 | 19.0 | 15.7 | 13.3 | 7.2 | 57.1 |
| 48 | 4.0 | 12.2 | 10.2 | 23.3 | 21.3 |
| 49 | 20.0 | 11.4 | 14.8 | 43.2 | 31.0 |
| 50 | 27.0 | 11.7 | 9.3 | 17.2 | 59.3 |
| **51** | **3.0** | **9.7** | **3.1** | **67.0** | **76.4** |
| 52 | 45.0 | 9.4 | 20.8 | 74.0 | 105.0 |
| 53 | 105.0 | 13.4 | 9.4 | 42.4 | 83.1 |
| 54 | 109.0 | 14.6 | 7.8 | 62.3 | 28.0 |
| 55 | 8.0 | 8.6 | 6.7 | 3.4 | 27.7 |
| **56** | **145.0** | **5.3** | **3.8** | **8.0** | **29.7** |
| 57 | 23.0 | 11.2 | 10.7 | 28.6 | 24.5 |
| 58 | 79.0 | 10.8 | 5.8 | 27.6 | 54.3 |
| 59 | 84.0 | 9.6 | 6.3 | 1.9 | 3.5 |
| 60 | 9.0 | 14.1 | 11.2 | 5.8 | 7.1 |
| CHARMM36m | | | | | |
| **1** | **9.0** | **5.9** | **3.1** | **6.1** | **25.7** |
| 2 | 102.0 | 16.4 | 10.7 | 27.5 | 3.9 |
| 3 | 30.0 | 8.5 | 12.7 | 16.2 | 90.5 |
| 4 | 92.0 | 11.0 | 12.2 | 35.4 | 64.7 |
| **5** | **30.0** | **8.2** | **3.1** | **12.3** | **73.0** |
| **6** | **167.0** | **8.3** | **4.1** | **6.7** | **20.8** |
| 7 | 15.0 | 10.7 | 7.4 | 12.8 | 66.0 |
| **8** | **65.0** | **10.0** | **3.6** | **10.6** | **44.9** |
| 9 | 14.0 | 8.7 | 4.7 | 20.6 | 25.9 |
| 10 | 43.0 | 8.0 | 8.4 | 15.9 | 10.2 |
| **11** | **240.0** | **5.1** | **3.4** | **5.0** | **41.8** |
| 12 | 17.0 | 13.7 | 7.9 | 68.1 | 26.8 |
| 13 | 79.0 | 6.0 | 11.1 | 49.7 | 52.1 |
| 14 | 27.0 | 8.2 | 6.8 | 16.3 | 7.2 |
| **15** | **54.0** | **10.2** | **3.6** | **47.9** | **46.2** |
| 16 | 20.0 | 12.2 | 6.5 | 49.8 | 31.6 |
| 17 | 23.0 | 9.5 | 7.6 | 6.3 | 18.7 |
| 18 | 126.0 | 21.8 | 14.0 | 16.3 | 16.3 |
| 19 | 6.0 | 8.9 | 10.4 | 12.5 | 57.3 |
| 20 | 2.0 | 16.4 | 8.2 | 90.0 | 26.0 |
| 21 | 3.0 | 13.4 | 12.7 | 32.6 | 28.2 |
| 22 | 45.0 | 8.2 | 8.0 | 26.2 | 9.8 |
| 23 | 149.0 | 11.7 | 6.7 | 21.3 | 76.7 |
| 24 | 87.0 | 15.8 | 10.4 | 15.8 | 80.1 |
| 25 | 86.0 | 11.4 | 7.2 | 34.6 | 58.2 |

| | | | | | |
|---|---|---|---|---|---|
| **26** | **38.0** | **7.0** | **3.2** | **20.8** | **45.6** |
| 27 | 77.0 | 15.6 | 9.0 | 24.8 | 36.0 |
| **28** | **161.0** | **10.8** | **3.5** | **5.2** | **63.8** |
| 29 | 72.0 | 12.1 | 10.1 | 24.7 | 29.0 |
| 30 | 8.0 | 18.2 | 24.3 | 2.7 | 19.6 |
| 31 | 41.0 | 9.9 | 13.8 | 18.8 | 5.2 |
| 32 | 14.0 | 11.4 | 8.6 | 16.9 | 45.1 |
| 33 | 63.0 | 11.8 | 9.0 | 38.3 | 109.1 |
| **34** | **111.0** | **10.6** | **3.1** | **28.1** | **37.8** |
| 35 | 64.0 | 13.1 | 8.0 | 101.1 | 5.4 |
| 36 | 9.0 | 8.5 | 8.0 | 29.2 | 22.1 |
| 37 | 152.0 | 11.7 | 8.4 | 1.0 | 32.1 |
| 38 | 160.0 | 13.8 | 8.9 | 49.5 | 33.2 |
| 39 | 216.0 | 11.7 | 10.5 | 23.6 | 35.9 |
| 40 | 82.0 | 10.4 | 10.4 | 17.0 | 47.4 |
| 41 | 7.0 | 11.8 | 13.2 | 35.6 | 74.1 |
| 42 | 12.0 | 18.6 | 7.4 | 70.9 | 66.1 |
| 43 | 74.0 | 16.8 | 8.9 | 66.3 | 109.4 |
| 44 | 86.0 | 20.2 | 10.4 | 78.1 | 29.2 |
| 45 | 25.0 | 10.5 | 9.4 | 6.9 | 33.6 |
| 46 | 3.0 | 15.5 | 15.1 | 15.1 | 6.4 |
| **47** | **20.0** | **8.5** | **3.1** | **68.2** | **13.3** |
| 48 | 23.0 | 14.2 | 7.0 | 42.1 | 64.4 |
| 49 | 16.0 | 5.1 | 8.5 | 36.2 | 19.0 |
| **50** | **50.0** | **8.7** | **3.7** | **11.5** | **55.3** |
| 51 | 68.0 | 14.3 | 9.3 | 41.5 | 9.3 |
| 52 | 24.0 | 11.5 | 12.2 | 25.8 | 40.0 |
| 53 | 121.0 | 9.0 | 9.2 | 47.9 | 26.3 |
| 54 | 149.0 | 11.6 | 7.8 | 11.6 | 82.1 |
| 55 | 21.0 | 13.3 | 8.9 | 24.8 | 64.5 |
| 56 | 19.0 | 9.4 | 10.6 | 20.7 | 39.3 |
| 57 | 6.0 | 18.7 | 7.8 | 73.3 | 59.2 |
| 58 | 182.0 | 10.8 | 9.5 | 15.7 | 1.3 |
| 59 | 10.0 | 15.6 | 8.8 | 20.9 | 1.8 |
| 60 | 2.0 | 18.3 | 9.1 | 48.6 | 11.8 |

Figure S3: E-SAM: S-SAM residue-residue contact map during the early times (< 50 ns) , intermediate (50-200 ns) and the final (> 200 ns) of protein-protein association in the simulations.

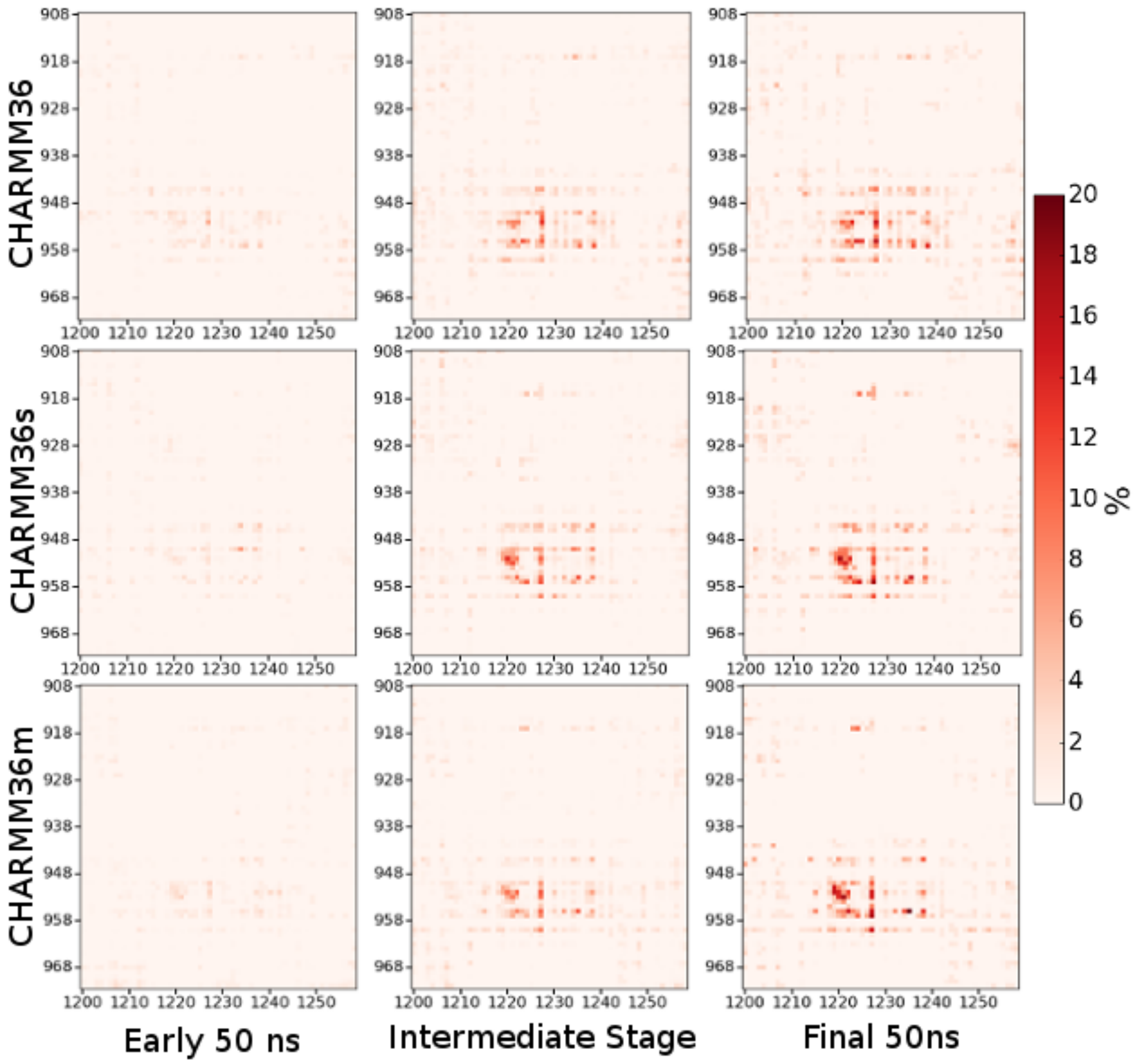

Figure S4: Time evolution of the number of contact, i-RMSD, buried solvent accessible surface area (SASA) and unscreened pair-interactions between E-SAM and S-SAM shown for a) simulation #2, run with CHARMM36s, b) #15 with CHARMM36m, c) #56 with CHARMM36s and d) simulation #1 with CHARMM36m.

a

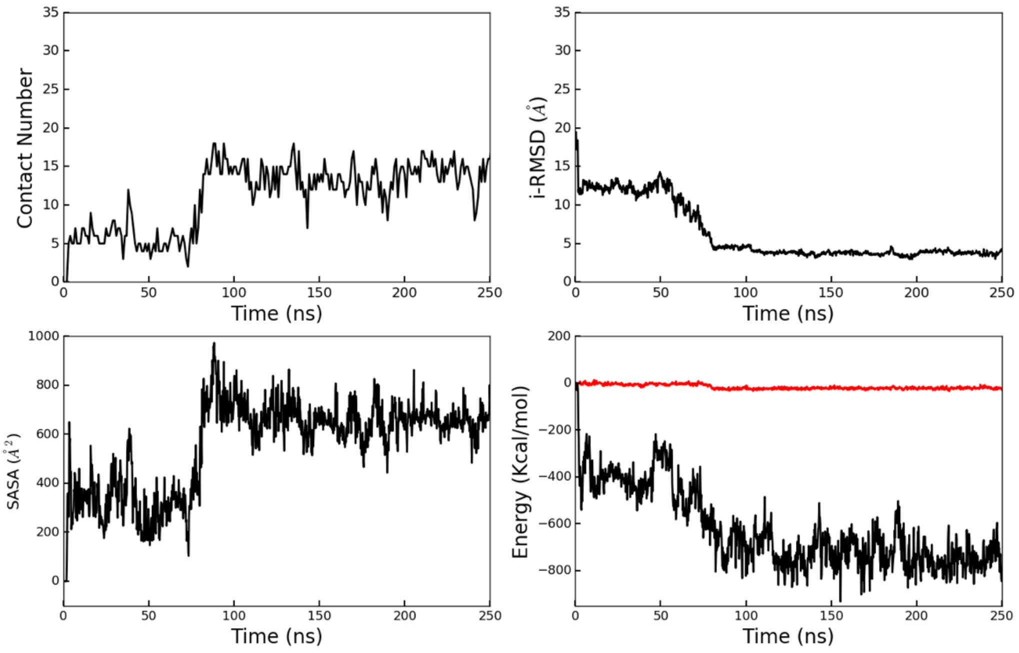


b

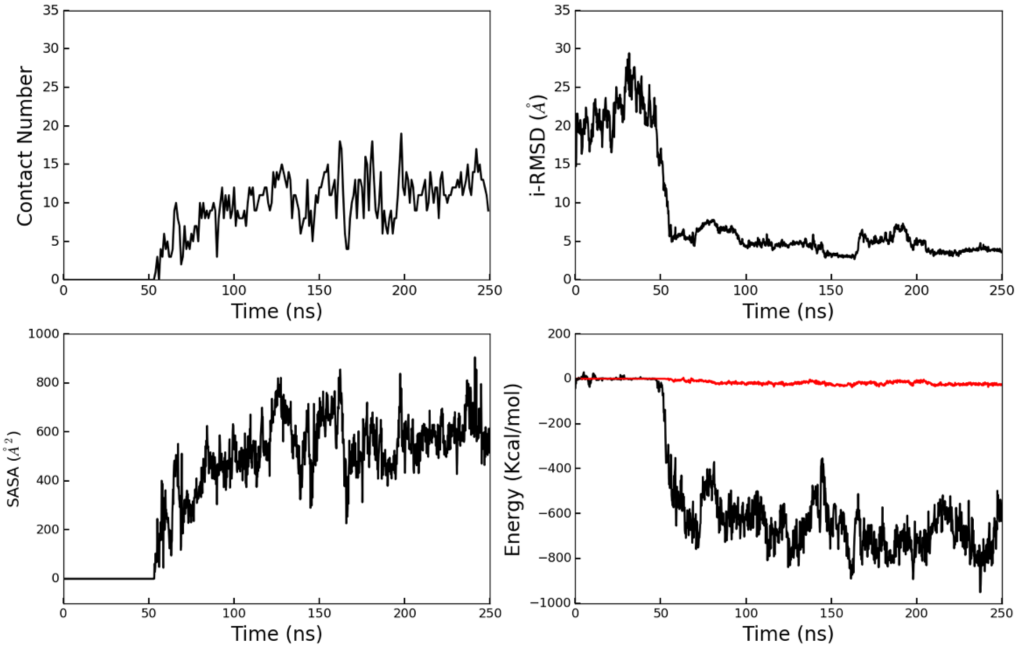

c

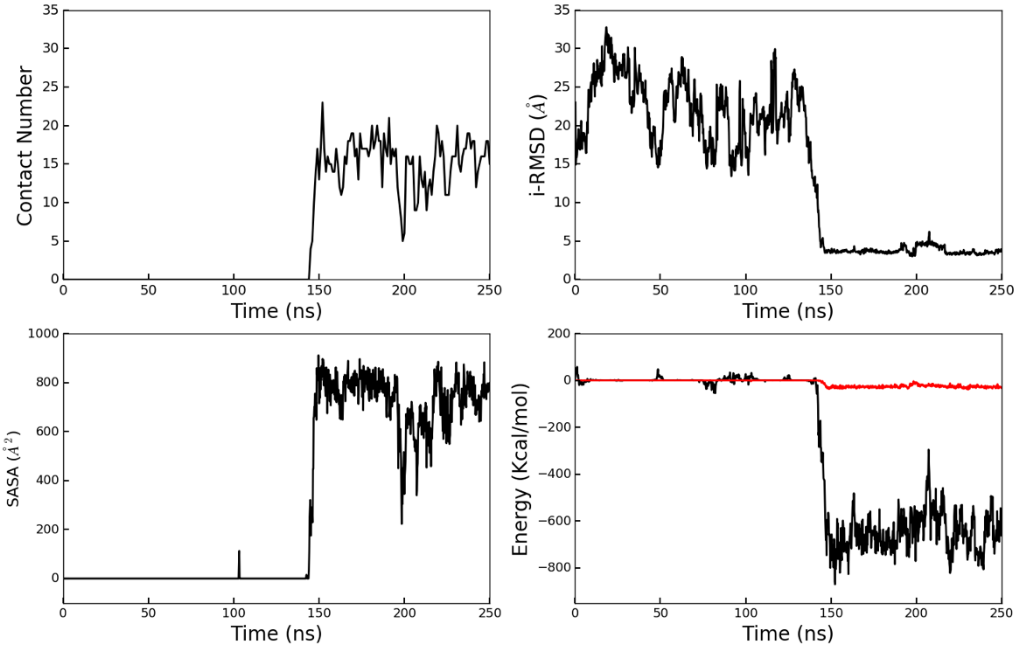

Contact Number
Time (ns)
i-RMSD (Å)
Time (ns)
SASA (Å²)
Time (ns)
Energy (Kcal/mol)
Time (ns)


d

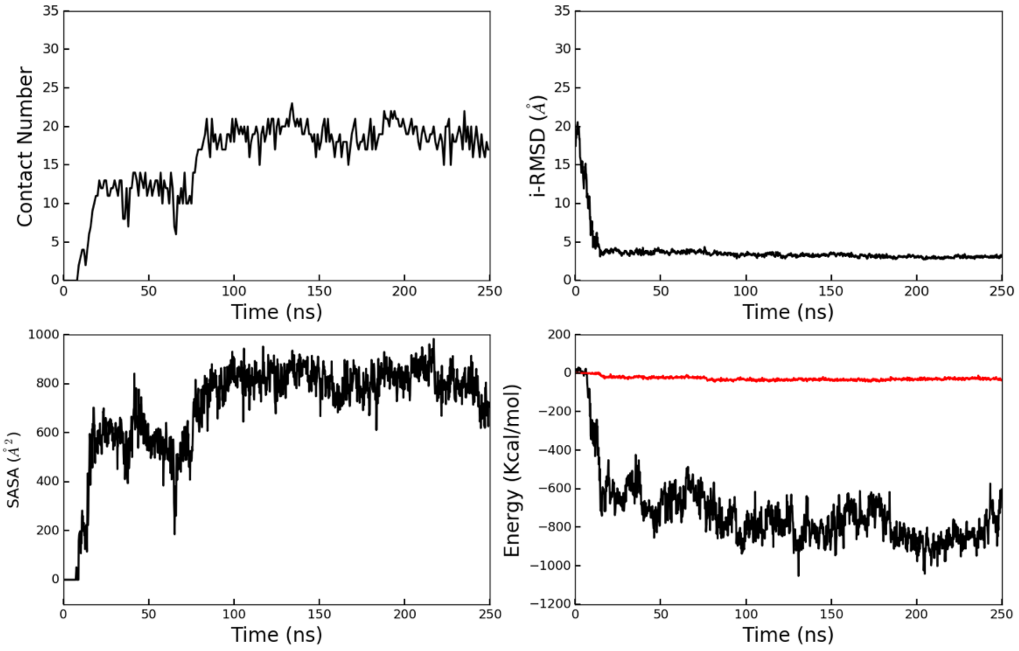

Contact Number
Time (ns)
i-RMSD (Å)
Time (ns)
SASA (Å²)
Time (ns)
Energy (Kcal/mol)
Time (ns)

Figure S5: Pair Interaction between E-SAM and S-SAM versus i-RMSD (Å) a) Analyzed for the last 50 ns of all trajectories and b) for the last 50ns of the trajectories that go to the native complex.

a

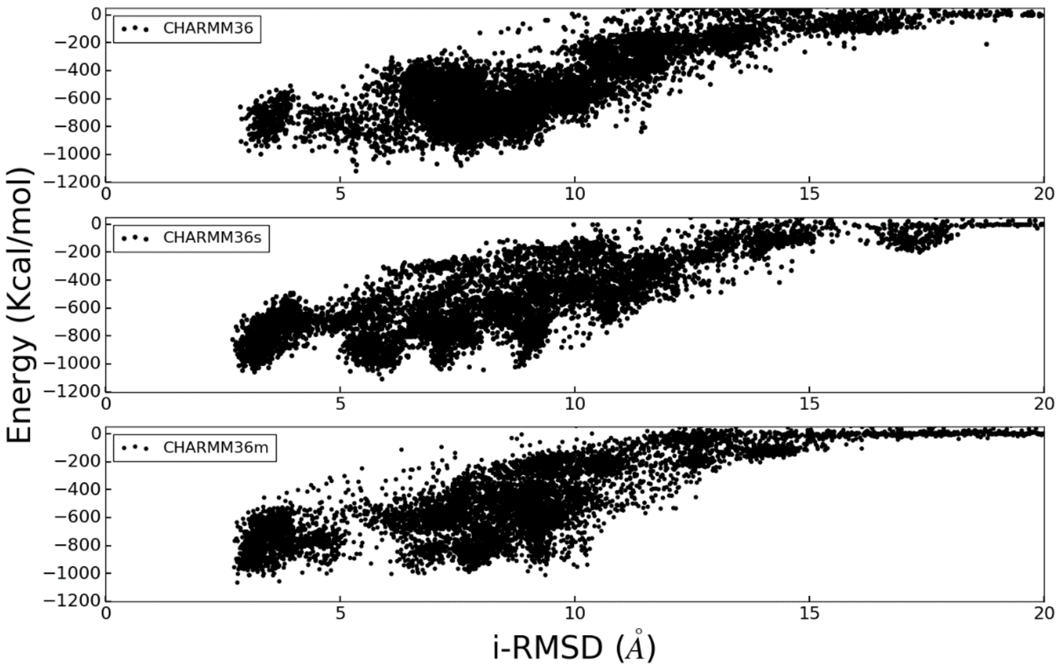


b

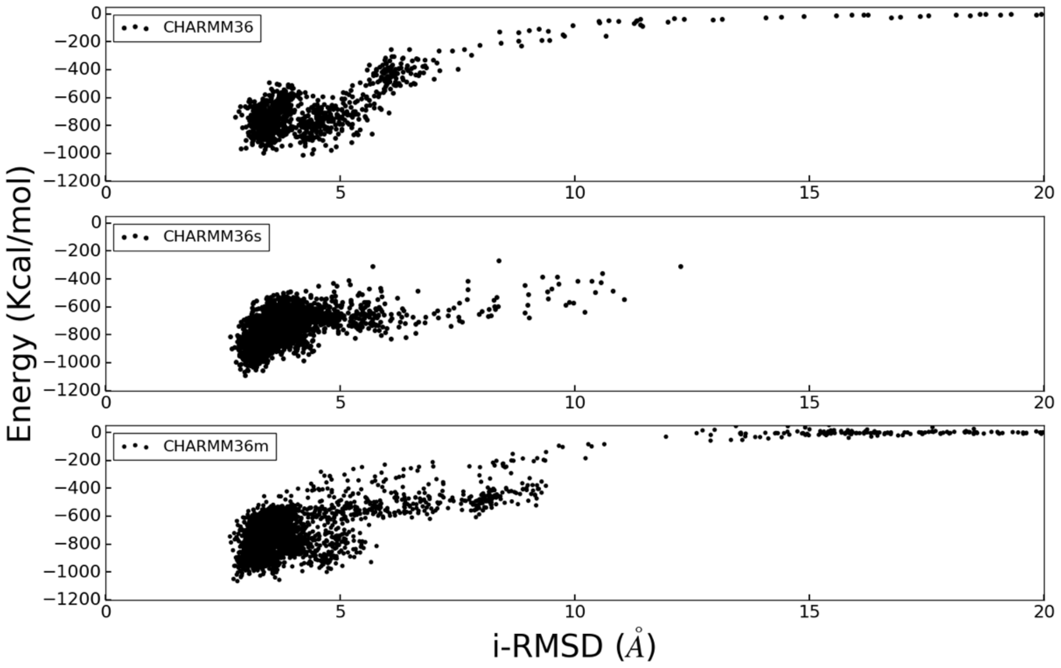
